# Epigenetic reprogramming driving successful and failed repair in acute kidney injury

**DOI:** 10.1101/2024.01.20.576421

**Authors:** Yoshiharu Muto, Eryn E. Dixon, Yasuhiro Yoshimura, Nicolas Ledru, Yuhei Kirita, Haojia Wu, Benjamin D. Humphreys

**Affiliations:** Division of Nephrology, Department of Medicine, Washington University in St. Louis, St. Louis, MO, USA; Department of Developmental Biology, Washington University in St. Louis, St. Louis, MO, USA

## Abstract

Acute kidney injury (AKI) causes epithelial damage followed by subsequent repair. While successful repair restores kidney function, this process is often incomplete and can lead to chronic kidney disease (CKD) in a process called failed repair. To better understand the epigenetic reprogramming driving this AKI-to-CKD transition we generated a single nucleus multiomic atlas for the full mouse AKI time course, consisting of ∼280,000 single nucleus transcriptomes and epigenomes. We reveal cell-specific dynamic alterations in gene regulatory landscapes reflecting especially activation of proinflammatory pathways. We further generated single nucleus multiomic data from four human AKI samples including validation by genome-wide identification of NF-kB binding sites. A regularized regression analysis identifies key regulators involved in both successful and failed repair cell fate, identifying the transcription factor CREB5 as a regulator of both successful and failed tubular repair that also drives proximal tubule cell proliferation after injury. Our interspecies multiomic approach provides a foundation to comprehensively understand cell states in AKI.

## Introduction

Acute kidney injury (AKI) is characterized by a sudden decrease in renal function, usually from ischemic or toxic insults(*1*, *2*). AKI is very common especially among hospitalized patients with an incidence up to ∼15%, and approaching ∼50% amongst critically-ill elderly patients(*2*). AKI causes short term morbidity and mortality, but it is also presents substantial risk of the development of future chronic kidney diseases (CKD), termed the AKI-to-CKD transition(*1–3*). Several lines of evidence have demonstrated that the AKI-to-CKD transition involves inflammation driving subsequent interstitial fibrosis(*3*). Tubular cell injury causes proinflammatory changes that promotes an innate immune response through recruitment and activation of immune cells, further promoting local inflammation(*3*). Both resident macrophages as well as infiltrating monocytes differentiate into distinct proinflammatory or profibrotic subsets depending on microenvironmental cues, and contribute to remodeling in the microenvironmental (*4*, *5*). Proinflammatory and profibrotic mediators activate other interstitial cell types, leading to loss of parenchyma and irreversible interstitial fibrosis, the final and common pathway in CKD(*6*). The molecular mechanisms underlying both successful kidney repair and the AKI-to-CKD transition remain incompletely understood. During AKI and subsequent repair, the epigenetic dynamics across diverse kidney cell types are especially poorly described since until recently methods to characterize these cell-specific changes were unavailable. Deciphering epigenetic changes during injury and repair will be important to identify new therapeutic targets for intervention(*7*).

We and others have generated single nucleus transcriptomic atlases from mouse kidneys after ischemia-reperfusion injury (IRI) and identified a *Vcam1*-expressing subset of proximal tubular cells (PTC) that emerges and persists after kidney injury(*8–10*). This cell state is characterized by a unique proinflammatory gene expression signature and it persists later after injury, leading us to name it failed-repair PTC (FR-PTC) (*8*). Furthermore, similar PTC subtype expressing VCAM1 was also observed in healthy and CKD human kidneys(*11–13*). The frequency of FR-PTC in advanced human CKD due to autosomal dominant polycystic kidney disease (ADPKD) was also markedly increased, replacing normal PTC(*12*). The increase in FR-PTC in CKD suggests a potential causative role in the AKI-to-CKD transition as well as CKD progression broadly. However, the molecular mechanisms driving failed repair gene expression signature as well as role in interstitial fibrosis driving CKD progression are incompletely understood.

Epigenomic alterations have gained attention recently as drivers of the AKI-to-CKD transition(*14*– *16*). Previously we have defined cell-type-specific epigenetic signatures in human healthy(*11*), ADPKD and CKD (*12*, *13*) kidneys, highlighting enrichment of NF-κB transcription factor binding sites on chromatin accessible regions in FR-PTC. We have interpreted increased chromatin accessibility for NF-κB binding motifs in the failed-repair PT state as reflecting a central role for this proinflammatory transcription factor in promoting inflammation, fibrosis and the AKI-to-CKD transition. Consistent with this, Gerhardt *et al.* recently performed a single-nucleus multiomic analysis on genetically-labelled *Ki67*+ cells at two timepoints after IRI (4 weeks, 6 months), describing proinflammatory transcriptomic and epigenetic alterations in *Vcam1*+ FR-PTC cells even 6 months after the insult(*17*), suggesting persistent epigenomic alterations after tubular injury. The lack of comprehensive, cell-type-specific epigenomic analysis along multiple time points from the acute to chronic phase of AKI hampers our understanding the epigenetic mechanisms driving the FR-PTC state.

Here, we performed single-nucleus ATAC-seq (snATAC-seq) on mouse kidneys following IRI to comprehensively describe the cell-specific epigenetic states and temporal dynamics in AKI-to-CKD transition. This single nucleus epigenetic atlas was integratively analyzed with single-nucleus RNA-seq (snRNA-seq) data from the same samples(*8*), and we describe the cis-regulatory epigenetic network driving failed repair cell states. To validate our findings for mouse kidneys with IRI, we further generated paired multimodal single nucleus multiomic datasets from four human AKI kidney samples. Interspecies multimodal analysis identified CREB5 as a critical transcription factor potentially involved in both the recovery and failed-repair processes in PT cells. We also generated an interactive data visualization tool for single nucleus epigenomic data (http://humphreyslab.com/SingleCell/). This detailed temporal single nucleus multiomic atlas mouse kidney IRI reveals comprehensive cell-specific epigenetic landscape during the IRI disease course, and provides a foundation to better understand disease mechanisms.

## Results

### Single nucleus chromatin accessibility atlas for mouse kidneys with ischemia-reperfusion injury

We performed snATAC-seq on a total of 19 male mouse kidney samples collected along six time points (sham, 4 and 12 h, and 2, 14 and 42 days after bilateral ischemia-reperfusion injury [IRI]; n = 3-4 per time point) with 10X Genomics Chromium Single Cell ATAC v1 (Fig.1A). The snRNA-seq for these samples was previously reported(*8*). After batch quality control (QC) filtering and preprocessing (See also Method), label transfer (Seurat) was performed on the snATAC-seq dataset using the snRNA-seq dataset(*18*, *19*). The snATAC-seq datasets were filtered using a 60% confidence threshold for cell type assignment to remove heterotypic doublets (Fig. S1A). Nuclei with inconsistent annotations between label transfer and manual annotation based on known cell type marker gene activities were further removed as remaining doublets and low-quality nuclei (Fig. S1B-F, See also Methods). The resultant datasets with high-quality nuclei were integrated with Harmony(*20*) and visualized in UMAP space (Fig. 1B). We detected 193,731 accessible chromatin regions among 157,000 nuclei in the final dataset. The differentially accessible regions (DAR) in each cluster includes the transcription start site (TSS) of known cell type marker genes (Fig. 1C), confirming our cell type annotations. Gene activities were predicted by accessibility of a gene body and promoter and used to further confirm our cell type annotations (Fig. S2). The QC metrics of snATAC-seq fragments in each time point indicated that the quality of samples during time course following ischemic injury were maintained (Fig. S3A-C). The QC metrics among cell types indicates that proximal convoluted and straight tubules (PCT/PST) have the best quality (Fig. S3D-F), although other cell types also showed sufficient quality to evaluate the differentially accessible regions (Fig. 1C) and transcription factor motif enrichment in accessible regions.

**Figure 1.**
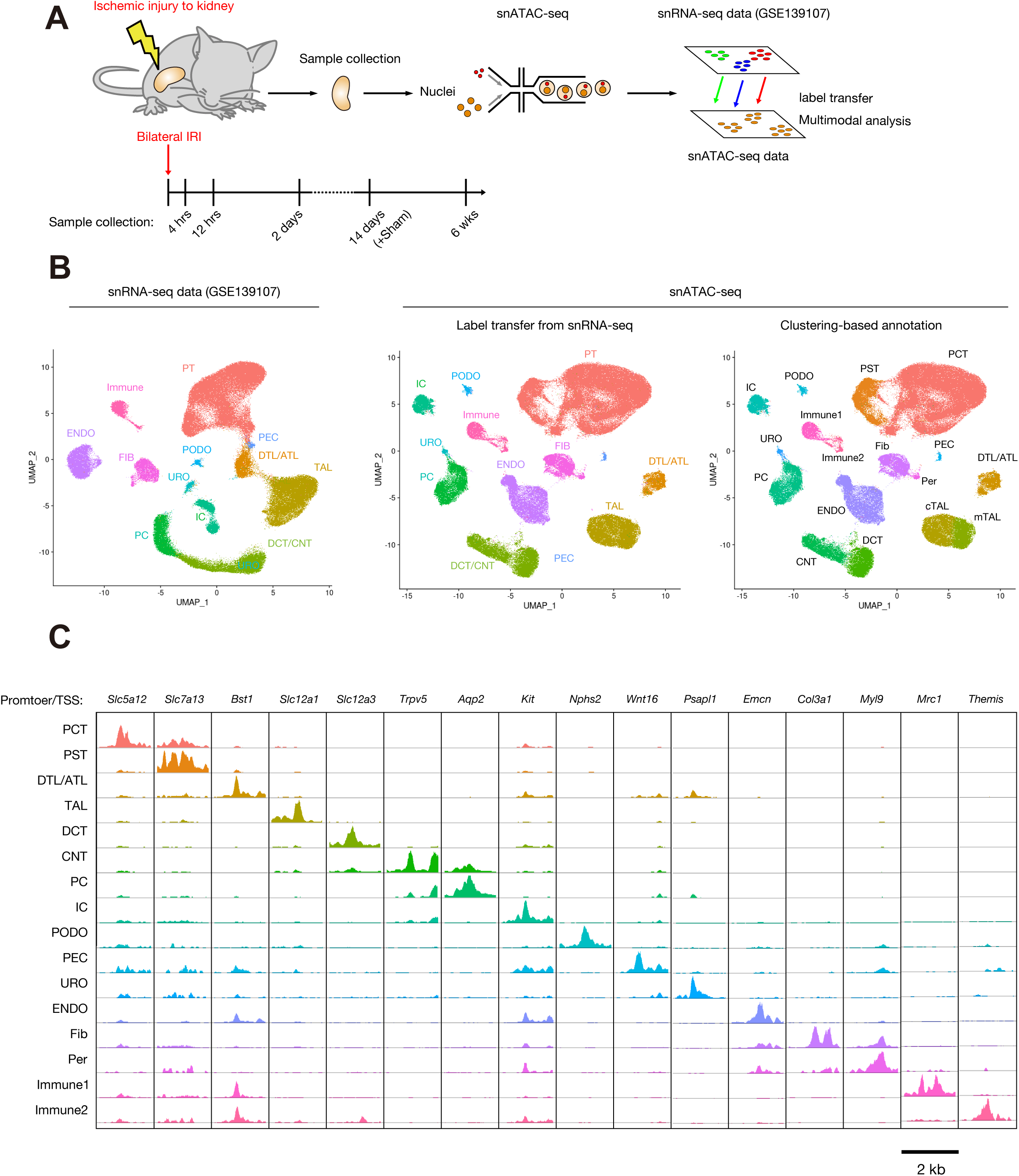
Single nucleus epigenetic profiling for mouse kidneys with IRI along time course. (**A**) Overview of experimental methodology. Single nucleus chromatin accessibility atlas was generated from mouse kidneys along time course after bilateral ischemia-reperfusion injury (IRI, n = 3-4 for each time point) and analyzed together with previously generated snRNA-seq dataset. (**B**) UMAP plot of previously generated snRNA-seq dataset (left) and newly sequenced snATAC-seq dataset with annotation by label transfer from snRNA-seq dataset (middle) and clustering-based annotation (right). PT, proximal tubule; PCT, proximal convoluted tubule; PST, proximal straight tubule; PEC, parietal epithelial cells; DTL/ATL, descending / ascending thin limb of Henle’s loop; cTAL/mTAL; cortical / medullary thick ascending limb of Henle’s loop; DCT, distal convoluted tubule; CNT, connecting tubule; PC, principle cells; ICA, Type A intercalated cells; ICB, Type B intercalated cells; PODO, podocytes; ENDO, endothelial cells; FIB (Fib), fibroblasts; Per, pericyte; Immune, immune cells; URO, uroepithelium. Clustering for snRNA-seq data was performed in our previous study by Kirita et al (*8*). (**C**) Fragment coverage (frequency of Tn5 insertion) around the differentially accessible regions (DAR) around each cell type at lineage marker gene transcription start sites. Scale bar indicates 2 kbp.

### Mouse proximal tubular cell heterogeneity after ischemia-reperfusion injury

We have previously identified failed-repair PT cells (FR-PTC) that emerge after IRI, adopt a proinflammatory and profibrotic phenotype, and persist in mouse kidneys(*8*). To better understand the epigenetic mechanism driving successful and failed repair of PT cells, we subclustered PTC/PST clusters (Fig.2A), identifying PT subtypes with mild to severe injury as well as the FR-PTC state which as expected had specifically increased accessibility to the *Vcam1* gene promoter (Fig. 2B). The injured PT subtypes display accessibility to the promoter of *Krt20*, previously found to be expressed in injured PT cells in mouse kidneys(*21*). Gene expression signatures in the snRNA-seq dataset(*8*) (Fig. S4A) were similar to gene activities predicted from accessibility of a promoter and gene body (Fig. S4B) among PT subtypes, confirming our PT subtype annotation. Transcription factor motif enrichment analysis with chromVAR (*22*) identified PT-subtype-specific transcription factor motif enrichment on open chromatin regions (Fig. 2C). The HNF4A binding motif was enriched in healthy PT subtypes and this was lost in injured and failed-repair cell states, consistent with previous findings in human kidneys(*11*). In contrast, motif enrichment for an AP-1 family transcription factor JUN/FOS as well as NFE2L2 were observed widely among injured PT states along the time course (Fig. 2D, Fig. S5). NFE2L2 regulates the expression of a cohort of antioxidant genes, protecting against oxidative stress induced by IRI and other renal insults(*23*). Interestingly, FR-PTC had enhanced NFE2L2 motif availability compared to healthy PT, suggesting a sustained oxidative stress response in FR-PTC (Fig. S5). The binding motifs for NF-κB family transcription factors were most enriched in FR-PTC (Fig. 2D), in agreement with previous findings in mouse and human kidneys(*8*, *11*, *17*). Together, these findings reveal distinct epigenetic programs between different PT cell states after injury, likely reflecting separate gene regulatory networks.

**Figure 2.**
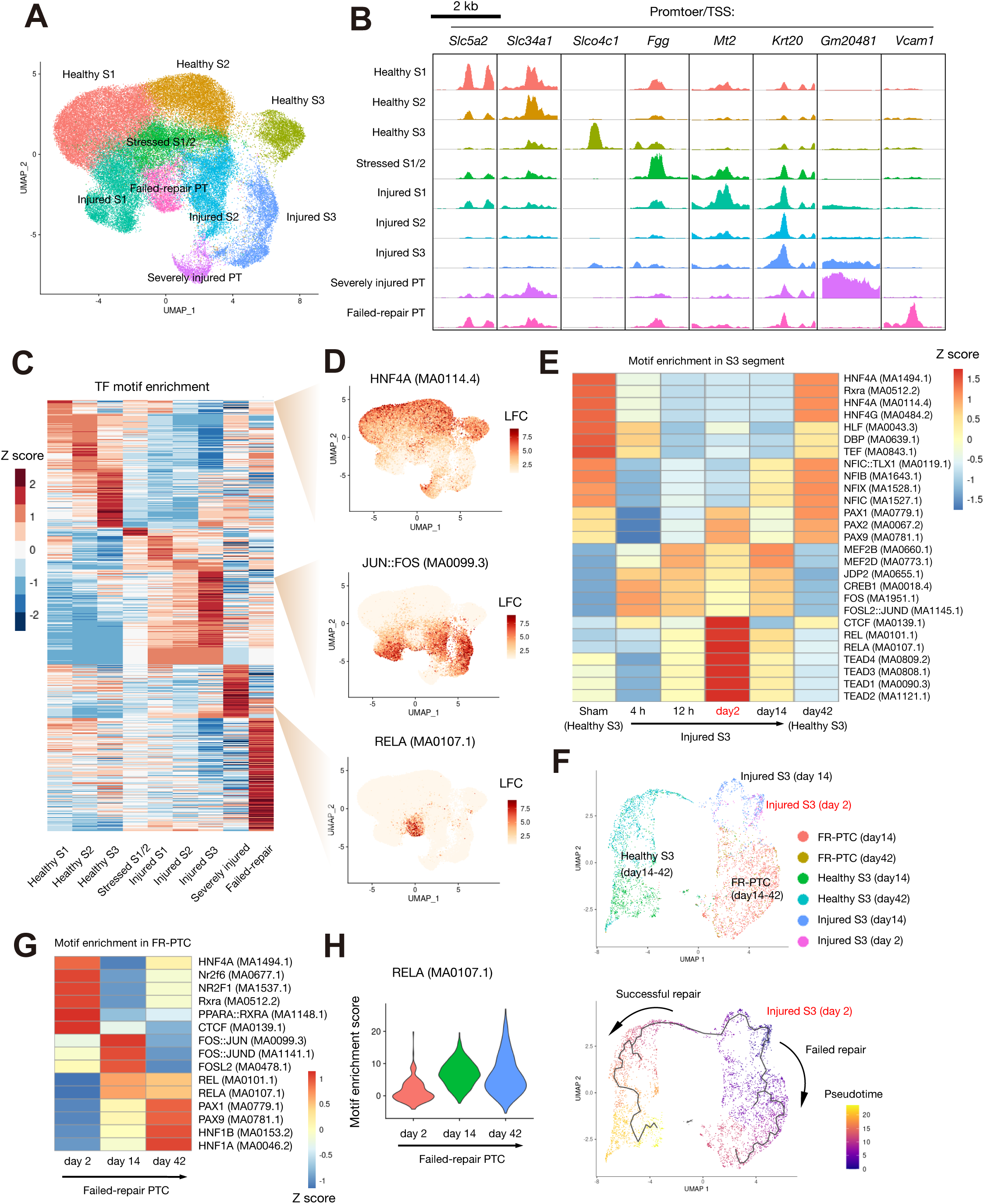
Heterogeneity of mouse proximal tubular cells following ischemia-reperfusion injury. (**A**) Subclustering of snATAC-seq PCT/PST on the UMAP plot with annotations by subtypes. (**B**) Fragment coverage around the TSS of differentially accessible regions (DAR) in each subtype. Scale bar indicates 2 kbp. (**C**) Heatmap showing relative transcription factor motif enrichment in each PT subtype. (**D**) UMAP plot showing chromVAR motif enrichment score in PT for HNF4A (MA0114.4, upper), JUN::FOS (MA0099.3, middle) and RELA (MA0107.1, low). The color scale for each plot represents a normalized log-fold-change (LFC) for the respective assay. (**E**) Heatmap showing relative motif enrichment for Healthy_S3 in sham and 42 days after ischemia-reperfusion injury (IRI), Injured_S3 at 4 h, 12 h, 2 days and 14 days after IRI. The most enriched motifs among injured S3 time points (4 h, 12h, 2 days and 14 days after IRI) were shown. (**F**) Pseudotemporal trajectory to model successful and failed recovery of injured PT S3 segment cells, constructed with injured S3 cluster at day2/day14, Healthy S3 cluster day14/day42 (successful recovery) and FR-PTC at day14/day42 (failed recovery) following IRI. Colored by the subtypes (upper) or pseudotime (lower). (**G**) Heatmap showing relative motif enrichment for FR-PTC at 2/14/42 days after IRI. The most enriched motifs in each time point were shown. (**H**) Violin plot displaying relative motif enrichment (chromVAR score) for RELA binding motif (MA0107.1) among FR-PTC at 2/14/42 days following IRI.

To further characterize epigenetic response following acute tubular injury, we evaluated the motif enrichment signature of the injured S3 segment over time (Fig. 2E)(*8*). Of note, in healthy S3 PTC, the transcription factor motif enrichment signatures were not completely recovered 6 weeks after IRI, consistent with the recent finding that transcriptional and epigenetic alteration in adaptive PTC persists for a long period, even 6 months after IRI(*17*). The S3 segment at day 2 was specifically enriched in NF-κB transcription factor motif availability as well as the Hippo pathway effector TEAD family molecules, and CCCTC-binding factor (CTCF) motif. CTCF is a multifunctional transcription factor serving as an architectural protein to create boundaries between topologically associating domains in chromosomes, regulating interactions between cis-regulatory regions and promoters(*24*). Previous studies have shown that ∼60% of CTCF-binding sites are cell-type-specific(*24*), suggesting that reorganization of CTCF availability may reflect global epigenomic remodeling in the injured S3 segment. To assess the cell fate of the injured S3 segment at day 2, we constructed a pseudotemporal trajectory (*25*) on the S3 segment population with injury (Injujred_S3: day2, 14) and recovery (Healthy_S3: day14, 42), and PT with failed recovery (FR-PTC: day14, 42) (Fig. 2F). Interestingly, injured S3 segment at day2 has two separate fates – either to healthy PT or to the failed-repair cell state. Given the transient activation of NF-κB binding motif in the injured S3 segment at day2 (Fig. 2E) and following its persistent activation in FR-PTC (Fig. 2D), NF-κB activation may initiate and promote the failed-repair branch of the trajectory (Fig. 2F).

NF-κB activation in FR-PTC was most prominent at day 14, and decreased at day 42 (Fig. 2G, H), although NF-κB activation was still prominent compared to other cell states at day 42 (Fig. S6). AP-1 transcription factor motif enrichment observed at day 14 was also diminished at day 42 (Fig. 2G), indicating resolution of acute inflammation. In contrast, PAX family transcription factor motif enrichment was increased in FR-PTC at day 42 compared to day 14 (Fig. 2G). Among PAX family genes, *Pax8* expression was relatively abundant in FR-PTC compared to healthy subtypes at day 42 after IRI in snRNA-seq (Fig. S7)), although *Pax8* expression level was highest among injured PT S3 segment in the whole dataset (Fig. S7A). PAX8 was recently found to promote renal cell carcinoma (*26*, *27*), which was also shown to differentiate from VCAM1+ PT cells (*28*). This finding suggests that PAX8 may contribute to a FR-PTC gene expression signature at the later stage of the time course after IRI. These analyses on PT cells collectively indicate that temporal dynamics of cell-type-specific transcription factor activation is associated with tubular injury and recovery after IRI and that NF-κB is potentially involved in the determination of PT cell fate after injury.

### Genome wide proximal tubule RELA binding sites

Our motif enrichment analysis on mouse PTC following IRI (Fig. 2) as well as previous findings(*8*, *11–13*, *17*) indicates activation of the NF-κB pathway in FR-PTC. The accessibility of transcription factor binding motifs allows for prediction of transcription factor activity(*22*), but it represents an inference and not an actual measurement of transcription factor-DNA interaction. To validate these predictions, we utilized cleavage under targets and release using nuclease (CUT&RUN) to directly identify genome-wide RELA binding sites in primary human PTC (*29*). We detected RELA binding to genomic DNA in primary human PTC after TNFα treatment (Fig. 3A), which activates the canonical NF-κB pathway(*30*). We identified 2,004 RELA binding peaks, which include both genomic regions with the H3K4me3+ promoter mark and those with H3K4me3-/ H3K27ac+ enhancer marks (Fig. 3A). Genes whose TSS was located close to RELA binding sites (within 2 kb from the TSS) were enriched with the genes of I-kappaB/ NF-kappaB complex (Fold enrichment: 64.55, FDR = 0.00088, Fig. 2B) among cellular component GO terms, and NIK-NF-kappaB signaling (Fold enrichment: 12.01, FDR = 0.00017) among biological process GO terms. The RELA binding peaks were located in promoter regions within 2kb from TSS (20.8%), introns (44.5%) and distal intergenic regions (28.7%, Fig. 3C), broadly covering both promoters and enhancers. Next, we asked which mouse FR-PTC DARs contain conserved RELA binding sites. We lifted human RELA binding peaks to mm10 mouse genome (minimum ratio of bases that must remap: 0.1, 1,438 peaks, recovery rate: 71.7%)(*31*) The lifted RELA binding sites were intersected with mouse FR-PTC DAR (Fig. 3D), identifying 103 conserved RELA binding sites in FR-PTC DAR. The nearby genes of those DAR were enriched with chemokine activity (fold enrichment 28.4, FDR=0.034) (Fig. 3E), suggesting that NF-κB activation in FR-PTC may promote local inflammation through chemokine secretion.

**Figure 3.**
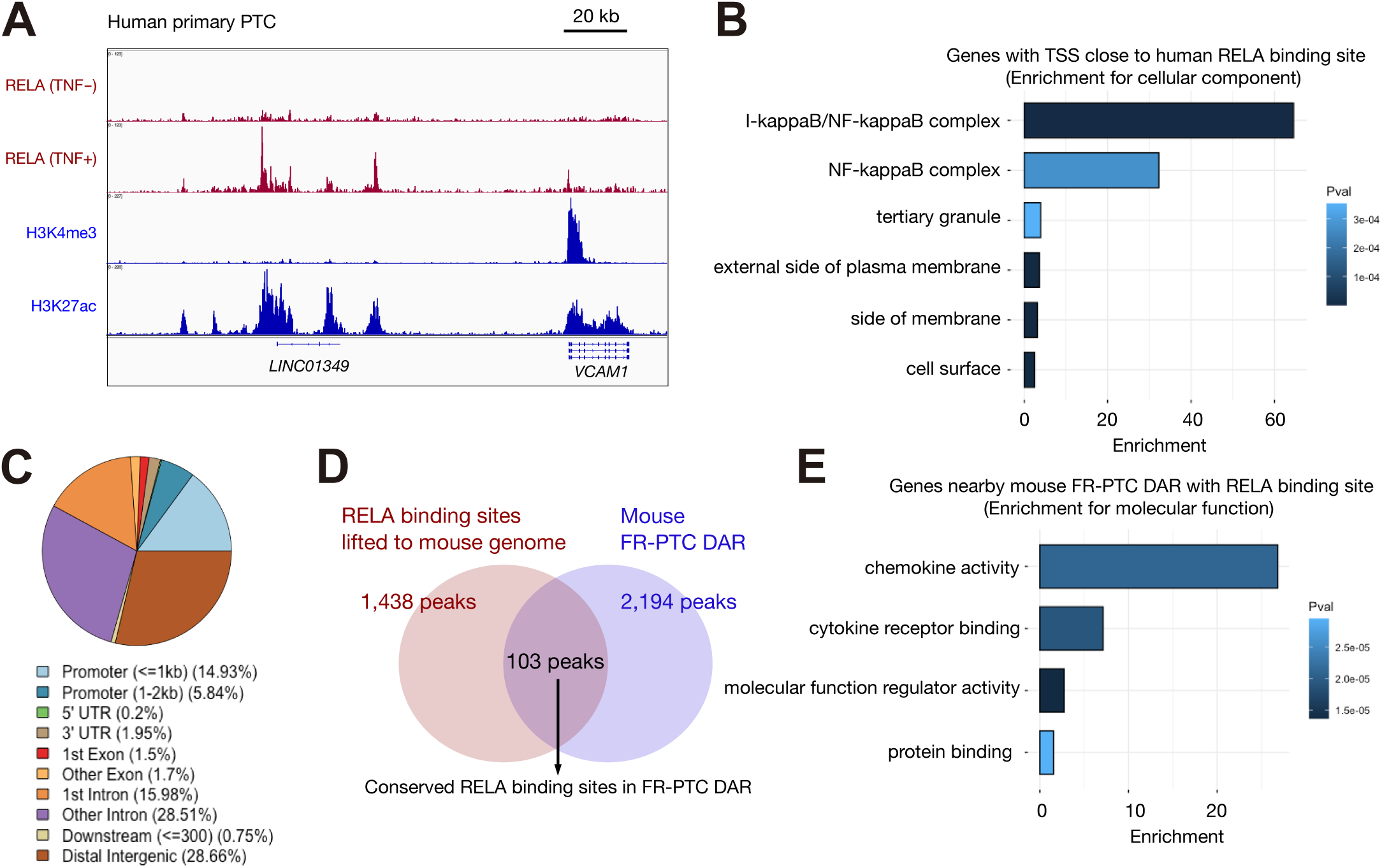
Genome wide proximal tubule RELA binding sites. (**A**) RELA binding sites with or without TNFα treatment (100 ng/ml, upper), and histone methylation and acetylation (lower) around *VCAM1* gene in human primary PTC determined by CUT&RUN assay. The scale bar indicates 20 kbp. (**B**) Cellular component gene ontology term enrichment analysis for the human genes with TSS close to RELA binding sites (<2 kbp). The six most enriched terms were shown. The color scale for each bar represents a P value. (**C**) Pie chart of annotated RELA binding site locations. (**D**) RELA binding peaks lifted to mm10 mouse genome were intersected with murine FR-PTC DAR, identifying 103 conserved RELA binding sites in FR-PTC DAR. (**E**) Molecular function gene ontology term enrichment analysis for the nearby mouse genes of conserved RELA binding sites in the murine FR-PTC DAR. The color scale for each bar represents a P value.

Among secretory molecules expressed in FR-PTC, *Ccl2* expression is prominently up-regulated (Fig. 4C)(*9*, *17*). CCL2 is a proinflammatory cytokine which has been implicated in AKI pathogenesis by recruiting immune cells to the local injured tubules(*32*, *33*). To characterize epigenetic mechanisms driving *Ccl2* up-regulation in FR-PTC, we constructed a cis-coaccessibility network(*25*) in FR-PTC around the *Ccl2* locus (Fig. 4A). A DAR with a conserved RELA binding site located at ∼2 kb upstream of *Ccl2* TSS was predicted to interact with the TSS, suggesting that this cis-regulatory region may activate *Ccl2* expression in a RELA-dependent manner in FR-PTC. Similarly, a total of five DAR with RELA binding sites were observed in the intron and 5’ distal region (within 100 kb) of the *Csf1* gene. Among them, the DAR at 30 kb upstream was predicted to strongly interact with the TSS of *Csf1* (Fig. 4D). *Csf1* encodes macrophage CSF (M-CSF or CSF1), a growth factor for monocyte / macrophage lineage and a major regulator of their proliferation and differentiation(*34*). Previous lines of evidence have shown the roles for CSF1 in tubule epithelial regeneration by expansion and M2 polarization of resident macrophages in the short term after renal injury(*35–37*), although in other contexts CSF1 expression can be detrimental, promoting tissue fibrosis(*38*, *39*). *Csf1* was up-regulated in injured PTC as well as FR-PTC (Fig. 4C) in agreement with the previous literatures (*35*, *37*, *37*). These results are consistent with the notion that RELA regulates expression of proinflammatory molecules through cis-regulatory regions.

**Figure 4.**
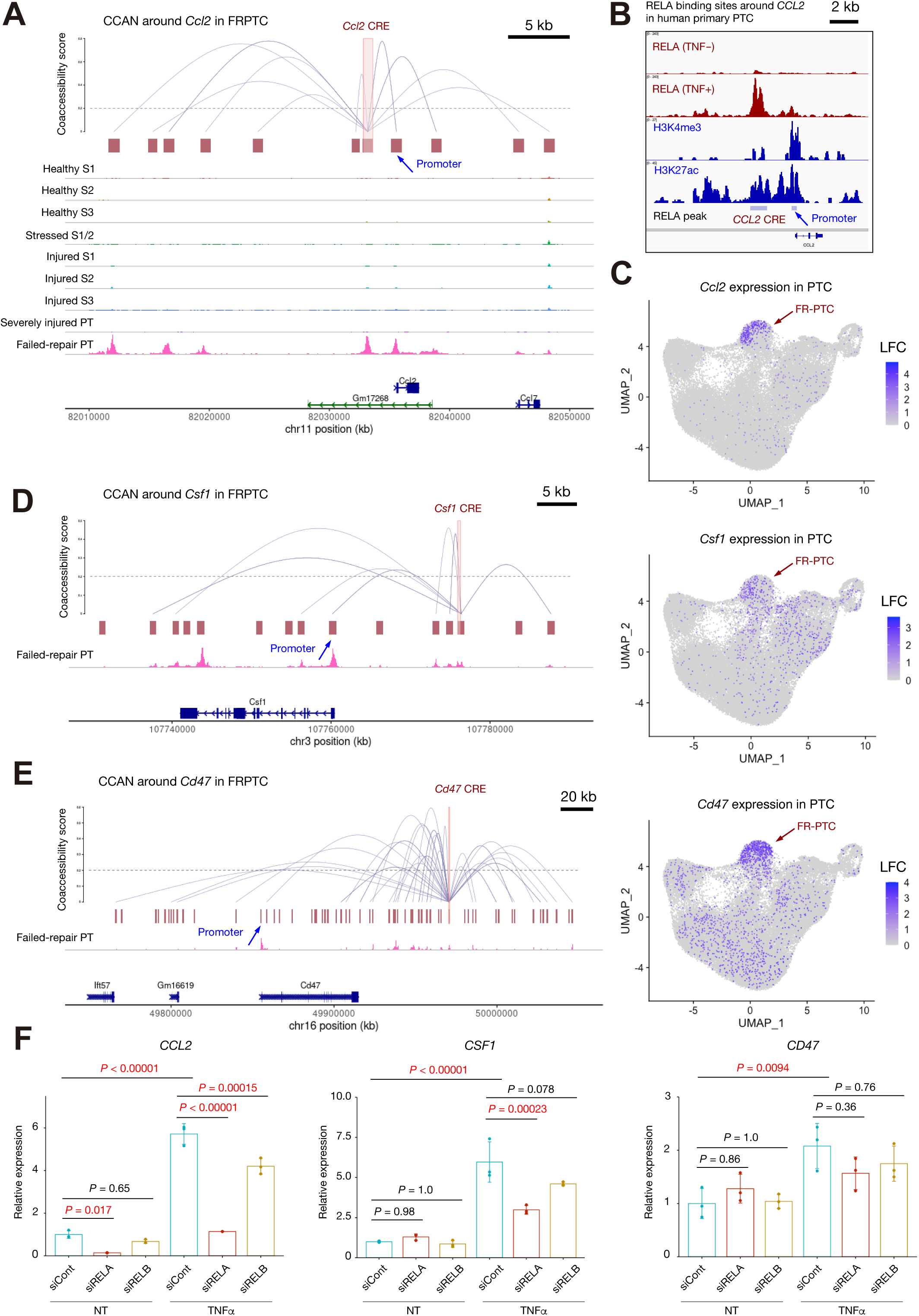
Cis-regulatory network driving inflammation by NF-κB signaling in failed-repair cells. (**A**) Cis-coaccessibility network (CCAN) predicts interactions (gray arcs) of a cis-regulatory region (CRE) with a conserved RELA binding site (chr11:82032734-82033526) and other accessible regions (red boxes) near the *Ccl2* gene in the mouse FR-PTC are shown (upper). Coverage plot showing accessible regions in each PT subtypes were also shown (lower). The scale bar indicates 5 kbp. (**B**) RELA binding site near *CCL2* gene identified by CUT&RUN assay for primary RPTEC treated with or without TNFα treatment (100 ng/ml, upper). CUT&RUN assay peaks for H3K4me3 and H3K27ac in primary RPTEC (middle), as well as RELA binding peaks (lower) were also shown. The scale bar indicates 2 kbp. (**C**) UMAP plot showing a gene expression level of *Ccl2* (upper), *Csf1*(middle) or *Cd47* (lower) among mouse PT subtypes. The color scale for each plot represents a normalized log-fold-change (LFC) for the respective assay. (**D, E**) The predicted interactions of a CRE with a conserved RELA binding site (chr3:107775891-107776242 (**D**) or chr16:49970506-49971086 (**E**)) and other accessible regions in the mouse FR-PTC (upper) around the *Csf1*(**D**) or *Cd47* (**E**) gene in the mouse kidneys. Chromatin accessibility in FR-PTC determined by snATAC-seq was also shown (lower). The scale bar indicates 5 kbp (**D**) or 20 kbp (**E**). (**F**) Quantitative PCR for *CCL2*, *CSF1* and *CD47* expression levels in primary human PTC with siRNA knockdown of *RELA* or *RELB* treated with or without TNFα (100 ng/ml) treatment. NT, no treatment. n = 3 biological replicates. Bar graphs represent the mean and error bars are the s.d. One-way ANOVA with post hoc Turkey test.

Genes upregulated in FR-PTC with conserved RELA binding sites in DAR also included non-secretory, membrane-bound signaling molecules involved in local inflammation. For example, *Cd47*, which encodes a membrane protein suppressing macrophage-mediated phagocytosis by sending a “*don’t eat me*” signal through interaction with signal regulatory protein alpha (SIRPα) (*40*). Indeed, *Cd47* expression was up-regulated in FR-PTC (Fig. 4C), suggesting a potential mechanism for the persistence of the FR-PTC for a prolonged period after the injury. The RELA-binding DAR located 3’ distal to *Cd47* gene was predicted to interact with accessible regions near the TSS (Fig. 4E).

We next investigated whether TNFα-induced expression of a subset of these genes was directly regulated by either RELA or RELB by siRNA knockdown in human PTC. Expression of these genes (CCL2, CSF1, CD47) predicted to interact with RELA in mouse FR-PTC were indeed up-regulated by TNFα treatment (Fig. 4F, Fig. S8), although the extent to which *RELA* contributes to their gene expression level was variable (Fig. 4F). *CCL2* expression was highly dependent on *RELA* regardless of TNFα treatment. The non-canonical NF-κB regulator *RELB* also regulated *CCL2* expression, although its regulation is more dependent on *RELA*. Of note, *RELB* expression was also regulated by *RELA* in TNFα-treated cells (Fig. S8). *CSF1* expression is also regulated by *RELA* in a TNFα-dependent manner. In contrast, the role of *RELA* in CD47 expression was limited in human primary PTC (Fig. 4F). Concurrent binding of other transcription may be needed for RELA to fully activate *CD47* expression. These findings collectively suggest that FR-PTC proinflammatory gene expression signature is shaped by numerous cis-regulatory regions, many but not all of which require NF-κB signaling

### Immune cells potentially regulated by failed-repair proximal tubular cells

Given epigenetic alterations driving expression of various cytokines and other membrane-bound signaling molecules interacting with immune cells in injured and failed-repair PTC (Fig. 4), transcriptional and epigenetic remodeling of immune cells may be coordinately regulated by PTC during AKI. Immune cells clustered into 6 cell types in our snRNA-seq dataset (Fig. 5A), as described in our previous work(*8*). Three subtypes are macrophages expressing *Mrc1* (Fig. 5A) which encodes a mannose receptor protein and marks alternatively activated macrophages (M2 macrophage) in both human and mice(*4*, *5*). We also observed *Flt3*-expressing dendritic cells (DC), T cells (Tcell) with *themis* expression and B cell (Bcell) with *Cd19* expression. Each of three *Mrc1*+ macrophage subtypes expressed unique markers (*Itga9*, *Plcb1*, *Gpnmb*). Among them, CCL2 receptor gene *Ccr2* was mainly expressed in *Plcb1*+ macrophage (*Plcb1*+Mac), and CSF1 receptor gene *Csf1r* was mainly detected in *Itga9*+ Mac, suggesting that FR-PTC may interact with heterogeneous macrophage subtypes (Fig. 5B) through distinct subsets of secretory molecules. Interestingly, *Sirpa* coding SIRPα protein, which receives the signal from CD47 to prevent phagocytosis, is expressed in *Plcb1+*Mac (Fig. S9). FR-PTCs recruit and activate *Plcb1+*Mac by CCL2, although they may inhibit their phagocytosis through CD47-SIRPα interaction.

**Figure 5.**
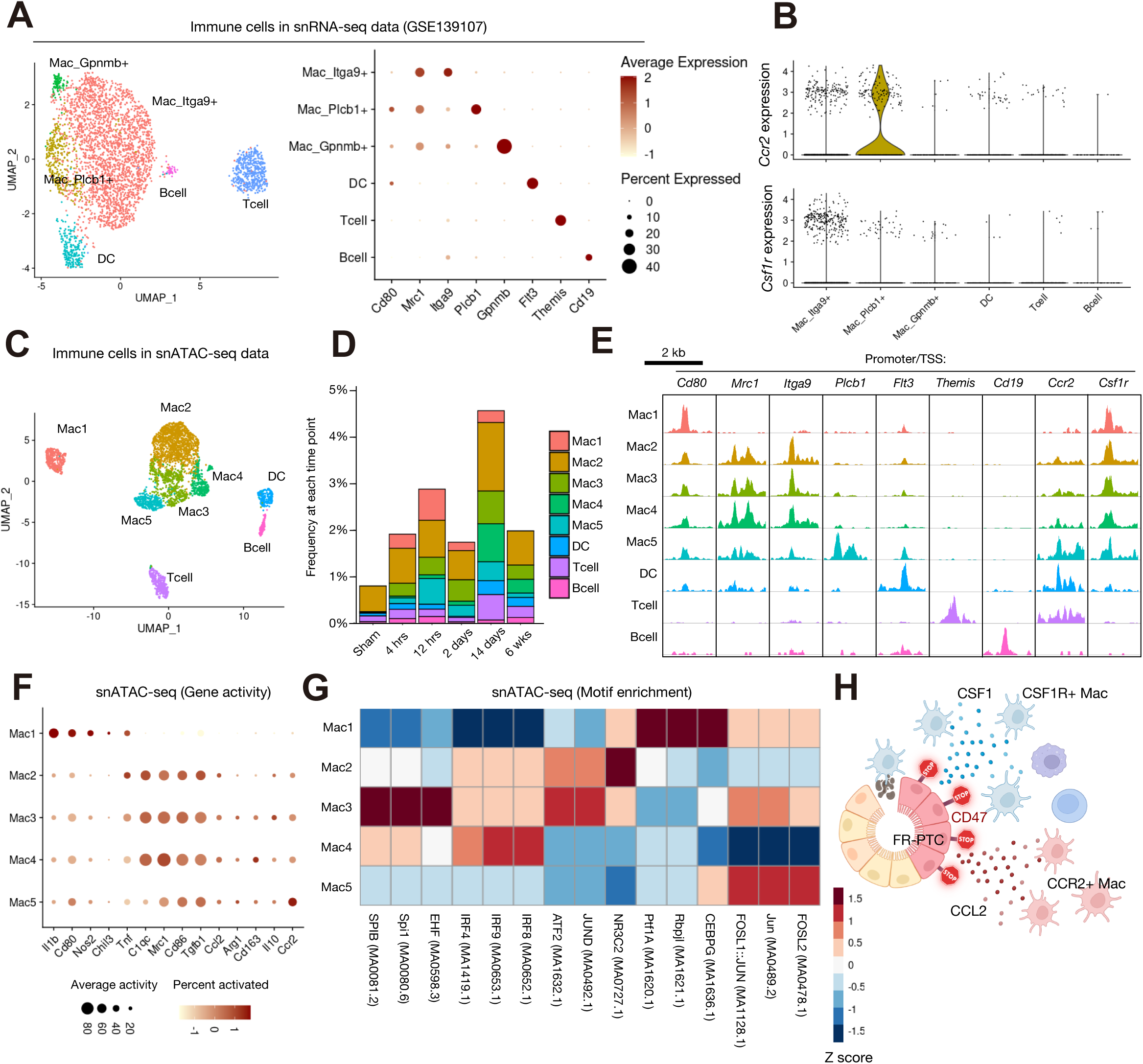
Immune cell landscape in mouse kidneys with ischemia-reperfusion injury. (**A**) Subclustering of mouse snRNA-seq immune cells on the UMAP plot (left) and dot plot showing expression patterns of the genes enriched in each of the subtypes (right). The diameter of the dot corresponds to the proportion of cells expressing the indicated gene and the density of the dot corresponds to average expression. Clustering was performed in our previous study by Kirita et al(*8*). (**B**) Violin plot showing gene expression level for *Ccr2* (upper) or *Csf1r* (lower) among immune cell subsets. (**C**) Subclustering of immune cells (Immune1, Immune2) on the UMAP plot of snATAC-seq dataset. (**D**) Frequency of each immune cell subpopulation for each time point in the snATAC-seq dataset. (**E**) Fragment coverage around the TSS of differentially accessible regions (DAR) in each subpopulation. Scale bar indicates 2 kbp. (**F**) Dot plot showing gene activities for the marker genes of macrophage polarization among macrophages in snATAC-seq data. (**G**) Heatmap showing relative transcription factor motif enrichment for macrophage subtypes. The most enriched motifs in each subtype were shown. (**H**) The proposed interaction of failed-repair proximal tubular cells (FR-PTC) and macrophages. While FR-PTC recruit and activate monocytes and macrophages through CCL2 and CSF1, they may escape from phagocytosis by CD47, leading to persistent inflammation. Schematic was created with BioRender.

To better understand macrophage heterogeneity in mouse kidney IRI, we subclustered immune cell clusters (Immune1, Immune2) from our snATAC-seq data, identifying 8 subtypes (Fig. 5C). The number and composition of these immune cell subtypes were dynamically increased immediately after IRI (Fig. 5D), and this increase persisted at 6 weeks after IRI, suggesting potential roles in the long-term tissue remodeling and AKI-to-CKD transition. Integration and label transfer from snRNA-seq data suggested robust identification of T cell, B cell, DC as well as *Itga9*+Mac (Fig. S10A, B). In contrast, *Plcb1*+Mac and *Gpnmb*+Mac prediction was weak, likely reflecting low numbers of immune cell nuclei. Nevertheless, the TSS of *Plcb1* was specifically accessible in Mac5 subtypes (Fig. 5E), suggesting that Mac5 is the *Plcb1*-expressing macrophage. Interestingly, we observed M1 macrophage with specific accessibility to *Cd80* TSS without *Mrc1* accessibility only in snATAC-seq (Fig. 5E), potentially due to low *Cd80* expression level. Mac2, 3, 4 subtypes displayed accessibility of the TSS in *Itga9* as well as *Mrc*1 gene, suggesting they are *Itga9*+ macrophage (Fig. 5E). The accessibility of *Ccr2* TSS was observed broadly among immune cells except B cell, although the accessibility was most significant in Mac4, consistent with high *Ccr2* expression level in *Plcb1*+Mac in snRNA-seq data. The gene activities predicted by snATAC-seq also suggested that Mac1 as M1 macrophage (*Il1b*, *Cd80*, *Nos2*) and that Mac2-5 were with M2 activation (*Mrc1*, *Cd86*, Fig. 5F). M2 macrophage was previously shown to contribute to tissue repair as well as tissue fibrosis through TGFβ signaling(*5*). Indeed, *Tgfb1* gene activity was predicted to be increased in Mac2-5 (Fig. 5F).

We next performed transcription factor motif enrichment analysis in macrophage subtypes (Fig. 5G). The resident macrophage subtype (Mac2) showed increased availability of mineral corticoid receptor NR3C2 binding motif compared to other macrophage subtypes (Fig. 5G) after renal injury (4 h and 2 days, Fig. S11C), consistent with previous literatures implicating mineral corticoid activity on the myeloid cells in renal fibrosis following kidney IRI(*41*, *42*). The most enriched motifs in *Ccr2*+ macrophage (Mac5) were AP-1 family transcription factors FOSL1 and JUN (Fig. 5G) that were activated immediately following IRI (Fig. S11D), likely reflecting numerous proinflammatory microenvironmental signals. These findings collectively suggest that FR-PTC interacts with various immune cell subsets by sending molecular cues, contributing to the macrophage heterogeneity following IRI (Fig. 5H).

### Proximal tubular cell heterogeneity in human acute kidney injury

To extend our multiomic atlas of mouse IRI, we next generated paired snRNA-seq and snATAC-seq datasets on the four kidney cortex samples of the patients with AKI (Table S1) and integrated these with previously published healthy kidney datasets (Fig. 6A)(*12*). Following low-QC nuclei filtering and preprocessing, cell types in the snRNA-seq dataset were identified by cell-type-specific marker gene expression (Fig. 6A, Fig. S12A). For snATAC-seq, we performed label transfer with Seurat using this snRNA-seq dataset (*18*, *19*). The snATAC-seq datasets were filtered using a 60% confidence threshold for cell type assignment. After remaining preprocessing (See also Methods, Fig. S13), integration of datasets with Harmony(*20*) and unsupervised clustering, we identified all the major cell types in snATAC-seq (Fig. 6B, Fig. S12B). The QC metrics both for snRNA-seq and snATAC-seq datasets was variable compared to control (Fig. S14), likely reflecting the fact that collection of human AKI samples was from postmortem samples compared to healthy samples that were from partial nephrectomies.

**Figure 6.**
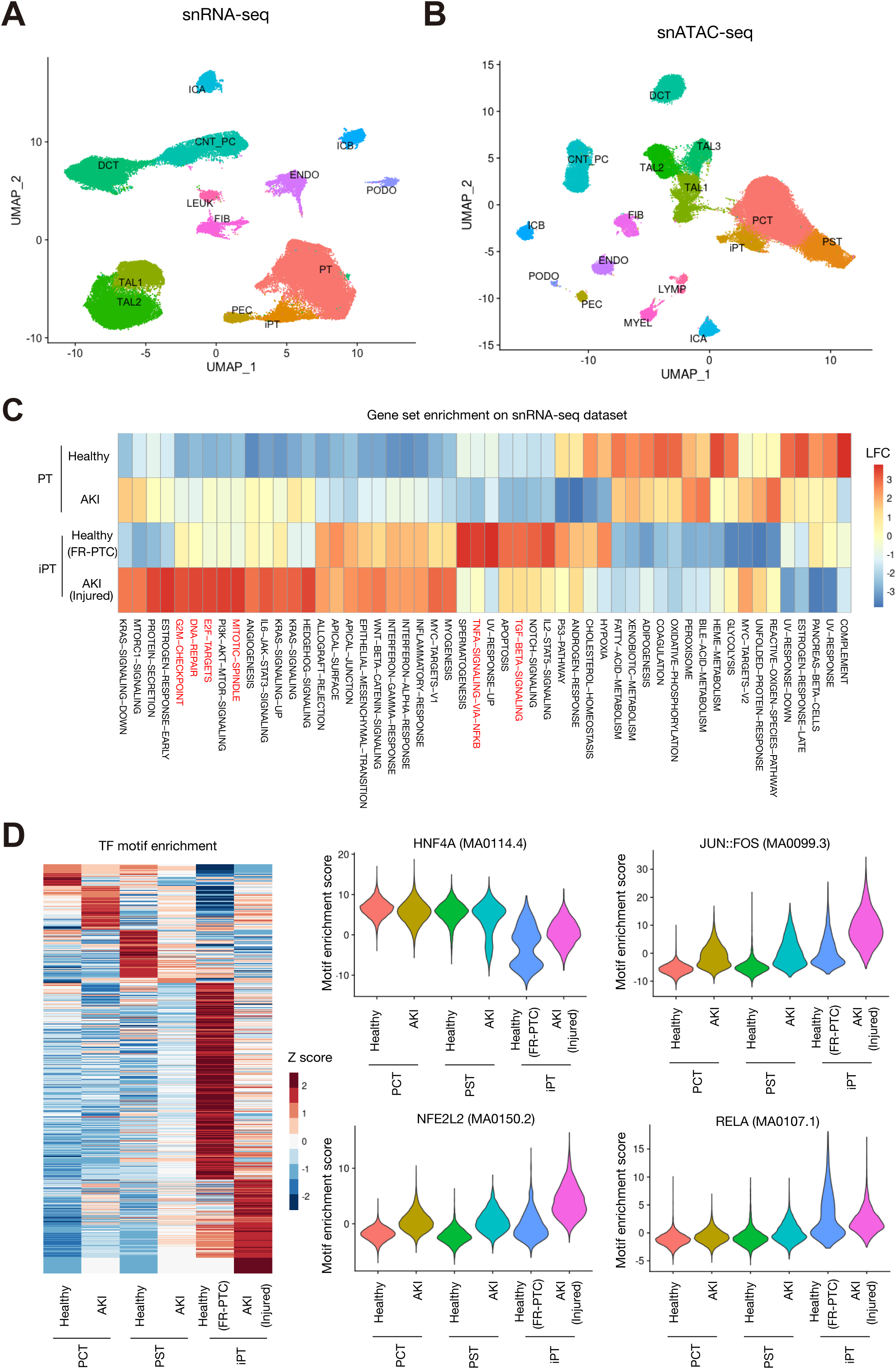
Heterogeneity of proximal tubular cells in the human kidneys with acute kidney injury. (**A, B**) UMAP plot of snRNA-seq (A) or snATAC-seq (B) data for human kidneys with acute kidney injury (AKI, n=4) or controls (n=5). (**C**) Heatmap showing relative enrichment of hallmark gene sets among PT lineages in snRNA-seq. (**D**) Heatmap showing relative motif enrichment among PT lineages in snATAC-seq (left). Violin plots displaying relative motif enrichment (chromVAR score) among PTC for HNF4A (MA0114.4, upper middle), NFE2L2 (MA0150.2, lower middle), JUN::FOS (MA0099.3, upper right) and RELA (MA0107.1, lower right).

*VCAM1*+PT (FR-PTC) in healthy kidneys and *HAVCR1*+ injured PTC in AKI kidneys were assigned to the same cluster after unsupervised clustering in both snRNA-seq (Fig.6A) and snATAC-seq (Fig. 6B), reflecting relative transcriptional and epigenetic similarities. We annotated this cluster as injury-related PT (iPT, Fig. 6A, B) containing both acutely-injured PT of AKI kidneys and failed-repair PTC of control kidneys. *VCAM1* expression was more abundant in iPT of control kidneys (FR-PTC) compared to iPT of AKI (acutely injured PT, Fig. S15A). The *VCAM1* promoter was accessible among iPT in both healthy and AKI kidneys, although it was more prominent among iPT of healthy kidneys (FR-PTC, Fig. S15B), consistent with transcriptional data (Fig. S15A).

Gene set enrichment analysis (*43*) with hallmark gene sets using the Molecular Signatures Database (MsigDB) suggested differential activation of molecular pathways between acutely injured PTC (iPT in AKI) and failed-repair PTC (iPT in control kidney) (Fig. 6C). The gene set associated with TNF signaling via NF-κB pathway was enriched in FR-PTC, consistent with specific enrichment of RELA binding motif in mouse FR-PTC (Fig. 2D). In contrast, the gene set related to proliferation such as G2/M checkpoint, E2F target and DNA repair gene set, reflecting proliferation and regeneration of proximal tubular cells in injured PTC. Transcription factor motif enrichment analysis for PTC in the snATAC-seq dataset (Fig. 6D) indicated HNF4A inactivation in both acutely injured PT and FR-PTC (Fig. 6D). JUN::FOS and NFE2L2 binding motifs were most enriched among acutely injured PTC (Fig. 6D). These findings were in agreement with the findings in mouse kidneys (Fig. 2D, Fig. S5). RELA motif enrichment was observed among both acutely injured PTC and FR-PTC (Fig. 6D), although gene set enrichment analysis indicated that the NF-κB target genes were more abundantly expressed in FR-PTC (Fig. 6C). Interestingly, we also observed transient enrichment of RELA binding motif among acutely injured PTC (day2) following mouse IRI (Fig. 2E). Increased RELA binding motif availability in acutely injured PTC may precede to the activation of RELA and subsequent up-regulation of its target genes in FR-PTC. Our multimodal single nucleus analysis in human AKI kidneys is limited because each is a snapshot in time. Nevertheless, our analyses of both human and mouse PTC collectively suggest that the PTC epigenetic alterations following AKI are largely conserved between human and mice.

### CREB5 drives proliferative recovery of proximal tubular cells after IRI

Next, we asked whether an interspecies approach might be useful to identify a novel molecular mechanism and potential therapeutic target in AKI. We recently developed RENIN (Regulatory Network Inference), a regularized regression approach to predict gene regulatory networks using multimodal single cell datasets (*44*). We applied RENIN to predict shared molecular mechanisms of PT cell state regulation shared between mouse and human. Several transcription factors were predicted to drive maladaptive cell states in human kidneys. *NFAT5*, *GLIS3*, *KLF6* and *CREB5* were identified as top-ranked accelerators of failed-repair cell states in both mouse and human kidneys (Fig. 7A). Interestingly, these transcription factors were also top-ranked as the transcription factors associated with injured cell states (Fig. 7A), suggesting that they are involved in both successful vs failed-repair in acute PT injury. *NFAT5* was previously described to be up-regulated following IRI(*9*) and predicted to promote failed-repair in healthy human kidneys using RENIN(*44*). *KLF6* was found to be induced after IRI, contributing to kidney injury in mice and humans(*45*). *GLIS3* is broadly expressed in renal tubules and localized in primary cilia. *GLIS3* deficiency was shown to induce polycystic kidney disease both in human and mice(*46*, *47*), although it has no known role in AKI. Of note, *PAX8* was top-ranked for promotion of both acute injury and failed repair in mouse PT. In contrast, *PAX8* was predicted to promote only the injured state but not the failed-repair state in human PT (Fig. 7A), indicating potential differences in PAX8 function during the AKI-to-CKD transition between human and mouse.

**Figure 7.**
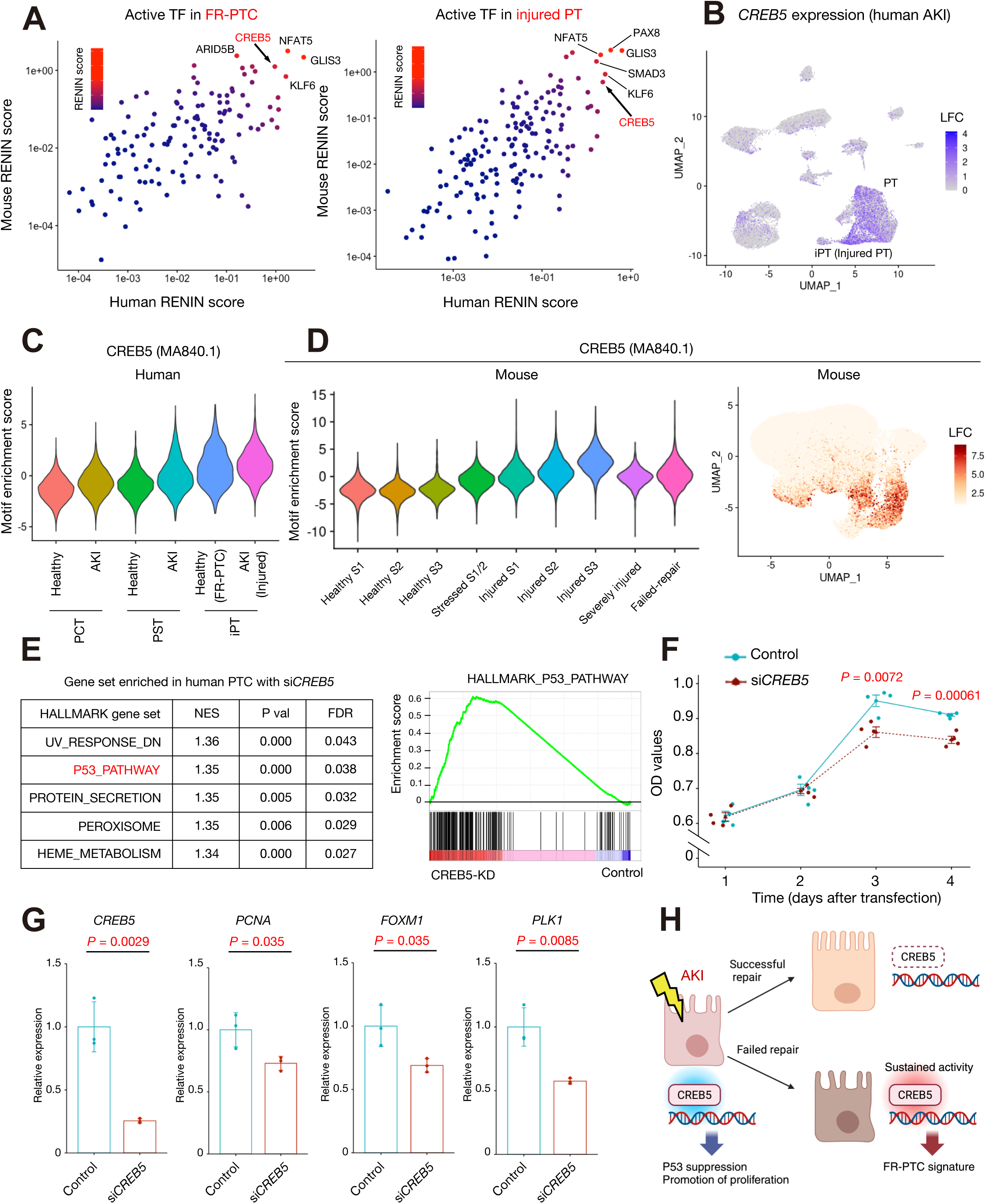
CREB5 drives proliferative recovery of proximal tubular cells after IRI. (**A**) The RENIN regulatory scores to predict transcription factor (TF) activities in the failed-repair proximal tubular cell (FR-PTC) state (left) or injured PTC state (right) compared to normal PT in human (x axis) and mouse (y axis) multiomics dataset. Higher regulatory score indicates more activation of the TF among failed-repair (left) or injured (right) cell state. (**B**) UMAP plot showing *CREB5* expression level in human AKI. The color scale represents a normalized log-fold-change (LFC). (**C**) Violin plot displaying relative motif enrichment (chromVAR score) among PT subtypes for CREB5 (MA0840.1) in human snATAC-seq dataset. (**D**) Violin plot (left) and UMAP plot (right) displaying relative motif enrichment among PT subtypes for CREB5 (MA0840.1) in mouse kidneys. The color scale for UMAP plot represents a normalized LFC. (**E**) Gene set enrichment analysis with the hallmark gene sets on published bulk RNA-seq data of primary human PTC with siRNA *CREB5* knockdown or control. The top-score gene sets with false discovery rate (FDR) < 0.05 (left) and enrichment of P53 pathway genes in human primary PTC with *CREB5* knockdown (right) were shown. NES, normalized enrichment score. (**F**) MTS assay in primary human PTC showing decreased proliferation by *CREB5* knockdown compared with control. Each time point (1-4 days after siRNA transfection) consists of n = 4 biological replicates. Student’s t-test for each time point. (**G**) Quantitative PCR for *CREB5*, *PCNA*, *FOXM1* and *PLK1* gene expression levels in primary PTC with *CREB5* knockdown compared with control (n = 3 biological replicates). Bar graphs represent the mean and error bars are the s.d. Student’s t-test. (**H**) The proposed role of CREB5 in PTC. Activation of CREB5 in acute injury may promote PTC proliferation and recovery from the injury. Persistent up-regulation of CREB5 in FR-PTC may contribute to FT-PTC gene expression signature. Schematic was created with BioRender.

*CREB5*, a member of the cAMP response element-binding protein family, was recently found to be highly expressed in various types of cancers including colorectal(*48*), ovarian(*49*), hepatocellular(*50*) and prostate cancers(*51*) and plays a promoting role in development of cancers. Moreover, a recent line of evidence suggested that CREB5 activation in PTC in the kidneys of the patients with diabetic kidney disease(*52*). *CREB5* expression was also up-regulated broadly in PTC in AKI (Fig. 7B, Fig. S16A-B), while its expression was localized in FR-PTC in control kidneys (Fig. S16A-B). In agreement with this finding, CREB5 motif activity was broadly increased in PTC in human AKI kidneys (Fig. 7C). In the mouse IRI dataset, CREB5 binding motif accessibility was enriched in FR-PTC as well as injured PTC, especially in the S3 segment, which is the most vulnerable site in hypoxic injury (Fig. 7D). Interestingly, up-regulation of *Creb5* expression was not clear in acutely injured mouse PTC (Fig. S16C), suggesting that enrichment of CREB5 binding motif in PT during acute phase of IRI may reflect translocation of preexisting CREB5 proteins to the nucleus from cytoplasm rather than up-regulation of CREB5 protein expression. In contrast, *Creb5* expression was prominently up-regulated in FR-PTC in snRNA-seq (Fig. S16C). To interrogate the role of CREB5 in PT recovery, we analyzed RNA-seq data of primary human PTC targeted CREB5 with siRNA knockdown. Gene set enrichment analysis implicated up-regulation of the P53 pathway by *CREB5* knockdown, which may inhibit proliferation and induce cell cycle arrest (Fig. 7E). Indeed, the primary human PTC with *CREB5* knockdown had slower proliferation compared to control (Fig. 7F), suggesting that CREB5 promotes cell proliferation and likely recovery in acutely injured PT states. This finding was consistent with down-regulation of proliferation marker *PCNA* as well as *FOXM1*; a master regulator of M phase progression, and its target *PLK1* by siRNA knockdown of *CREB5* in primary human PTC (Fig. 7G). These findings collectively suggest that CREB5 has a beneficial role in the recovery from acute injury by promoting proliferation, although persistent activation of CREB5 in FR-PTC may contribute to its unique gene expression signature (Fig. 7H). Together, these findings demonstrate the usefulness of interspecies multimodal approach to interrogate molecular mechanism of AKI and identify novel therapeutic targets.

## Discussion

We generated and analyzed single nucleus chromatin accessibility profiles across the spectrum of mouse acute injury and repair and compared to human AKI. Our integrative analysis of this atlas with our previously generated single nucleus transcriptomic atlas from the same samples(*8*) allows us to understand the temporal dynamics of gene regulation during injury, repair and failed repair. We elucidated cell-type-specific transcription factor activities with an emphasis on the NF-κB effector RELA in injured and failed-repair PTC states along time course. We mapped conserved RELA binding sites onto the mouse FR-PTC-specific accessible regions, revealing the cis-regulatory landscape driving failed-repair defining genes. Furthermore, we generated paired snRNA-seq and snATAC-seq datasets from human AKI kidneys and compared with mouse IRI, shedding light on conserved and divergent gene regulatory networks between human and murine AKI.

Given that various renal insults drive an increase of *VCAM1*-expressing FR-PTC along with infiltrating immune cells in both mouse and human kidneys(*8*, *10*, *12*, *13*, *17*), understanding the mechanism of emergence and maintenance of FR-PTC, and their biological roles will be critical to dissect the molecular mechanism driving AKI and AKI-to-CKD transition. Here we confirm and extend evidence implicating NF-κB activity as an important putative mediator of this cell state. Previous studies in mice have shown that suppression of NF-κB pathway reduced tubular apoptosis and chemokine-induced immune cell infiltration after IRI(*53*). Furthermore, phagocytosis of apoptotic cells by renal epithelial cells mediated by HAVCR1 recruits p85 and inactivates NF-κB, suppressing inflammatory pathway and protecting from IRI(*54*). These findings suggest that proximal tubule NF-κB activity may promote inflammation and exacerbate tubular injury during AKI. We confirmed enriched NF-κB motif availability among FR-PTC in mouse kidneys with IRI (Fig.2F), consistent with previous studies (*8*, *9*, *11*, *17*). Furthermore, the NF-κB pathway was activated in injured PTC as early as two days after injury before the FR-PTC increased (Fig. 2E). We identified mouse FR-PTC DAR with conserved RELA binding sites identified by CUT&RUN on primary human PTC(Fig.3D). The nearby genes of those DAR with conserved RELA binding sites were enriched with the genes for chemokines (Fig. 3E), that may allow FR-PTC to interact with surrounding other cell types, including immune cells. For instances, we demonstrated cis-coaccessibility network connecting conserved NF-κB binding DAR and TSS of signaling molecules like *Ccl2*, *Csf1* and *Cd47* as the potential modulators of local immune cell landscape (Fig.4, 5). Our findings highlight that NF-κB activation in FR-PTC may have a role in shaping local immune cell landscape during AKI-to-CKD transition (Fig. 5H).

Injury-induced expression of proinflammatory mediators in PTC includes CCL2, which was previously shown to promote inflammation and interstitial fibrosis after ischemic renal injury(*32*, *33*). *Ccl2* expression was up-regulated in FR-PTC likely in a RELA-dependent manner (Fig. 4). We identified myeloid lineage subsets including five macrophage subtypes as well as T cell and B cell after subclustering of immune cells in our dataset. Among them, *Plcb1+*Mac (Mac5) displayed the highest mRNA expression level as well as chromatin accessibility around TSS of the *Ccr2* gene, which encodes the CCL2 receptor. The number of the macrophages was increased and maintained at least until 6 weeks after IRI (Fig. 5D). In contrast, *Itga9+Mac* (Mac2) subtype exists in the mouse kidneys without IRI (Fig. 5D), suggesting that they are resident immune cells in the kidney. All the macrophage subtypes express *Csf1r* coding for the CSF1 receptor, that was also up-regulated in FR-PTC. CSF1 may be also stimulatory signal from PTC to activate macrophages in IRI. Interestingly, FR-PTC also expresses the gene encoding CD47,which protects cells from being phagocytosed by immune cells and is often overexpressed in various cancers to avoid phagocytosis by tumor-associated macrophages(*40*). CD47 is a ligand for SIRPα on macrophages, and binding CD47 to SIRPα initiates a molecular signaling to inhibit phagocytosis. Accumulated MRC1+ M2 macrophage initiates tissue fibrosis through TGFB1 expression, irreversible hallmark of CKD(*4*). These findings suggest that altered immune landscape by FR-PTC contributes to AKI-to-CKD transition.

To leverage our mouse dataset to identify a novel mechanism involved in AKI-to-CKD transition, we also generated single nucleus multimodal datasets for human AKI and intersected the gene regulatory networks between humans and mice. The gene regulatory network was constructed with RENIN, which we recently developed as a regularized regression approach (*44*) for multiomics datasets. Most of the transcription factors that were predicted to contribute to the injured state were also top-ranked for promoting factors of failed-repair state, suggesting considerable overlap of the transcription factors being activated between acutely injured and failed-repair PTC (Fig. 7A). Such transcription factors included CREB5, whose motif activity as well as mRNA expression were up-regulated in PTC in human AKI (Fig. 7). *CREB5* knockdown by siRNA in primary human PTC up-regulated the genes associated with P53 pathway, which may induce cell cycle arrest. Consistent with activation of P53 pathway, *CREB5* knockdown decreased proliferation of primary human PTC (Fig. 7E), suggesting a pro-proliferation and pro-recovery role for CREB5 after PTC injury. Interestingly, we also found *CREB5* knockdown partly reduced the failed-repair gene expression signature(*44*), implicating sustained activation of CREB5 may also contribute to the failed-repair state in AKI-to-CKD transition.

In summary, we performed multimodal single nucleus analysis of a mouse kidney IRI timecourse to define cell-type-specific, temporally dynamic gene regulations. We also generated human AKI multiomics datasets to validate our mouse findings as well as leverage our mouse time-course dataset to understand shared AKI mechanism and identify potential therapeutic targets. Our study is limited by the lack of spatial information of transcriptomic and epigenomic alteration. In the future, application of spatially resolved transcriptomics to the samples of kidneys with IRI will contribute to further dissection of intercellular communication in the AKI-to-CKD transition. Our single nucleus multimodal analysis of mouse IRI kidneys provides a foundation on which to base future efforts to develop better diagnostic and therapeutic approaches for AKI.

## Materials and Methods

### Experimental design

To comprehensively describe the cell-specific epigenomic perturbation in human and mouse AKI, we generated and sequenced snATAC-seq libraries from the frozen mouse kidney samples previously sequenced for snRNA-seq and published(*8*). Briefly, bilateral ischemia-reperfusion injury (IRI) on the kidney (18 min) was performed on male mice at 8-10 weeks of age, and control mice underwent sham surgery. Mice were euthanized with designated time point and the kidneys were collected and frozen with liquid nitrogen. All mouse samples were harvested according to the animal experimental guidelines issued by the Animal Care and Use Committee at Washington University in St. Louis. C57BL/6J were purchased from The Jackson Laboratory (Bar Harbor, ME).

Human AKI kidney cortical samples were obtained according to the protocol by the Washington University Institutional Review Board. Kidneys were discarded human donor kidneys from deceased patients (two male and two female, 55-69 years old, table S1). Samples were cut out from outer cortex and frozen in liquid nitrogen. The single cell dataset generated from control kidneys (10X Genomics Chromium Single Cell 3’ v3 chemistry and 10X Genomics Chromium Single Cell ATAC v1) were already published(*11*).

### Nuclear dissociation for library preparation

For snATAC-seq, nuclei were isolated with Nuclei EZ Lysis buffer (NUC-101; Sigma-Aldrich) supplemented with protease inhibitor (5892791001; Roche). Samples were cut into < 1 mm pieces, homogenized using a Dounce homogenizer (885302–0002; Kimble Chase) in 2 ml of ice-cold Nuclei EZ Lysis buffer, and incubated on ice for 5 min with an additional 2 ml of lysis buffer. The homogenate was filtered through a 40-µm cell strainer (43–50040–51; pluriSelect) and centrifuged at 500 g for 5 min at 4°C. The pellet was resuspended, washed with 4 ml of buffer, and incubated on ice for 5 min. Following centrifugation, the pellet was resuspended in Nuclei Buffer (10× Genomics, PN-2000153), filtered through a 5-µm cell strainer (43-50005-03, pluriSelect), and counted. For snRNA-seq preparation, the RNase inhibitors (Promega, N2615 and Life Technologies, AM2696) were added to the lysis buffer, and the pellet was ultimately resuspended in nuclei suspension buffer (1x PBS, 1% bovine serum albumin, 0.1% RNase inhibitor). All these processes were performed in a cold room at 4°C. Subsequently, 10X Chromium libraries were prepared according to manufacturer protocol.

### Single nucleus ATAC sequencing for mouse kidneys and bioinformatics workflow

For mouse IRI kidneys, the snATAC-seq libraries were generated using 10X Genomics Chromium Single Cell ATAC v1 chemistry following nuclear dissociation. Libraries were sequenced on an Illumina Novaseq instrument and counted with CellRanger ATAC v2.0 (10X Genomics) using mm10. The read configuration was 2x150bp paired-end. Sample index PCR was performed at 9 cycles. A mean of 457,383,804 reads were sequenced for each snATAC library (s.d.= 111,816,895) with a median of 22,026 fragments per cell (s.d.= 15,328). The data were aggregated with CellRanger ATAC v2.0, and the aggregated dataset (filtered_peak_bc_matrix) was processed with Seurat v4.0.2 and its companion package Signac v1.4.0(*55*). Low-quality cells were removed from the aggregated snATAC-seq library (subset the high-quality nuclei with peak region fragments > 2000, peak region fragments < 100000, %reads in peaks > 25, blacklist ratio < 0.08, nucleosome signal < 4 & TSS enrichment > 2). The barcodes representing doublets were determined with AMULET v1.1 (*56*) run on each snATAC-seq library, and those were filtered out from the integrated dataset (Fig.S1A). A gene activity matrix was constructed by counting ATAC peaks within the gene body and 2kb upstream of the transcriptional start site using protein-coding genes annotated in the Ensembl database (*19*). FindTransferAnchors and TransferData functions were used for label transfer from snRNA-seq dataset using gene activities, according to instructions(*55*). After label transfer, the snATAC-seq datasets were filtered using an 60% confidence threshold for low-resolution cell type assignment to remove low-quality nuclei and heterotypic doublets. Latent semantic indexing was performed with term-frequency inverse-document-frequency (TFIDF) followed by singular value decomposition (SVD). A KNN graph was constructed to cluster cells with the Louvain algorithm. Batch effect was corrected with Harmony(*20*) using the RunHarmony function in Seurat. After clustering and cell type annotation based on lineage-specific gene activity (Fig. S1B-D), the nuclei with inconsistency between predicted low-resolution cell type and annotation based on lineage-specific gene activity were further filtered out to remove remaining doublets and low-quality nuclei (Fig. S1C, E). After filtering out these artifacts, he dataset was processed for batch effect correction with Harmony(*20*), clustering and cell type annotation based on lineage-specific gene activity (Fig. 1). The final snATAC-seq library contained a total of 193,731 peak regions among 157,000 nuclei. The number of fragments in peaks per nucleus was a mean of 9292 +/- 6066, and %Fragments in peaks per nucleus was a mean of 53.1 +/- 12.7. Fraction of reads in peaks, number of reads in peaks per cell and ratio of reads in genomic blacklist regions per cell for each patient were shown in Fig. S3. Differential chromatin accessibility among cell types was assessed with the Seurat FindMarkers function for peaks detected in at least 10% of cells with a likelihood ratio test and a log-fold-change threshold of 0.25 to identify differential chromatin accessibility. The nearby genes were determined by ClosestFeature function. Bonferroni-adjusted p-values were used to determine significance at an FDR < 0.05.

For subclustering of PTC (PCT and PST) or immune cells (Immune1 and Immune2) in snATAC-seq data, the target cell types were extracted from the integrated dataset (Fig. 1B). Following TFIDF/SD and subclustering, the clusters with marker gene activities for other cell types were removed as remaining low QC nuclei and doublets. Subsequently TFIDF/SVD, processing with Harmony and subclustering was performed with UMAP (Fig. 2 and 5).

### Single nucleus RNA sequencing for human kidneys and bioinformatics workflow

For human AKI kidneys, snRNA-seq libraries were generated with 10X Genomics Chromium Single Cell 3’ v3 chemistry following nuclear dissociation. A target of 10,000 nuclei were loaded onto each lane. The cDNA for snRNA libraries was amplified for 15 cycles. Libraries were sequenced on an Illumina Novaseq instrument and counted with CellRanger v6.0.0 with --include-introns argument using GRCh38. The read configuration for the libraries was 2x150bp paired-end. A mean of 503,452,820 reads (s.d. = 18,797,210) were sequenced for each snRNA library corresponding to a mean of 54,336 reads per cell (s.d.= 26,822). The mean sequencing saturation was 38.8 +/- 8.0%. The mean fraction of reads with a valid barcode (fraction of reads in cells) was 39.6 +/- 15.4%.

The output of CellRanger (filtered_gene_bc_matrix) were processed through Seurat v4.0.2(*18*). Ambient RNA contamination was corrected for each dataset by SoupX v1.5.0(*57*) with automatically calculated contamination fraction. Each of datasets was then processed to remove low-quality nuclei (nuclei with top 5% and bottom 1% in the distribution of feature count and RNA count, and those with %Mitochondrial genes > 0.25). Heterotypic doublets were identified with DoubletFinder v2.0.3(*58*) assuming 8% of barcodes represent heterotypic doublets), and resultant estimated doublets were removed after merging datasets. The datasets were integrated in Seurat using the IntegrateData function with anchors identified by FindIntegrationAnchors function. Subsequently, the doublets and low QC clusters were removed for these datasets. The major cell types were identified in the dataset for AKI kidneys (Fig. S17). The control datasets (n=5) were previously published (GSE185948)(*12*). The AKI and control datasets were integrated with batch effect correction with Harmony v1.0(*20*) using RunHarmony function on assay RNA in Seurat.

Then, there was a mean of 8127+/- 1692 nuclei in control or 8058 +/- 2971 nuclei in AKI per snRNA-seq library. The number of unique molecular identifiers (UMI) per nucleus was a mean of 3536 +/- 1914 in control or 3906 +/- 2429 in AKI. The number of detected genes per nucleus was a mean of 2222 +/- 803 genes in control or 2434 +/- 864 genes in AKI. %Mitochondrial genes was 0.027 +/- 0.050% in control or 0.028 +/- 0.048% in AKI (Fig. S12). Clustering was performed by constructing a KNN graph and applying the Louvain algorithm. Dimensional reduction was performed with UMAP and individual clusters were annotated based on expression of lineage-specific markers (Fig. S13). Differential expressed genes among cell types were assessed with the Seurat FindMarkers function for transcripts detected in at least 10% of cells using a log-fold-change threshold of 0.25. Bonferroni-adjusted p-values were used to determine significance at an FDR < 0.05.

### Single nucleus ATAC sequencing for human kidneys and bioinformatics workflow

For human AKI kidney, snATAC-seq libraries were generated using 10X Genomics Chromium Single Cell ATAC v1 chemistry following nuclear dissociation. Libraries were sequenced on an Illumina Novaseq instrument and counted with CellRanger ATAC v2.0 (10X Genomics) using GRCh38. The read configuration was 2x150bp paired-end. Sample index PCR was performed at 9 cycles. A mean of 402,063,865 reads were sequenced for each snATAC library (s.d.= 54,544,038) with to a median of 16,333 fragments per cell (s.d.= 2,033). Five control snATAC-seq libraries (Control 1-5) were prepared and published in a prior study (GSE151302)(*11*). The libraries from control and AKI kidneys were aggregated with CellRanger ATAC v2.0 Subsequently, the aggregated dataset (filtered_peak_bc_matrix) was processed with Seurat v4.0.2 and its companion package Signac v1.4.0(*55*). Low-quality cells were removed from the aggregated snATAC-seq library (subset the high-quality nuclei with peak region fragments > 2500, peak region fragments < 25000, %reads in peaks > 15, blacklist ratio < 0.1, nucleosome signal < 4 & TSS enrichment > 2). A gene activity matrix was constructed by counting ATAC peaks within the gene body and 2kb upstream of the transcriptional start site using protein-coding genes(*19*) and used for label transfer with FindTransferAnchors and TransferData functions using snRNA-seq dataset for the same human AKI samples, according to instructions(*19*). After label transfer, the snATAC-seq datasets were filtered using an 50% confidence threshold for low-resolution cell type assignment to remove low-quality cells and heterotypic doublets. A low-quality cluster with simultaneous marker gene activities with multiple cell types was further removed. Following TFIDF/SVD, batch effect was corrected with Harmony(*20*). After clustering and cell type annotation based on lineage-specific gene activity (Fig. S14A-D), the nuclei with inconsistency between predicted cell type and annotation based on lineage-specific gene activity were further filtered out to remove remaining doublets and low-quality nuclei (Fig. S14E). The barcodes representing doublets determined with AMULET v1.1 (*56*) in each library were further filtered out from the integrated dataset (Fig. S14A). After filtering out these artifacts, dataset was processed with Harmony(*20*) to remove batch effect. Clustering and cell type annotation were based on lineage-specific gene activity (Fig. S13B). The final snATAC-seq library contained a total of 249,164 peak regions among 64,350 nuclei (31,397 nuclei for control and 32,953 nuclei for AKI) and represented all major cell types within the kidney cortex (Fig. 6B). The number of fragments in peaks per nucleus was a mean of 8629 +/- 3575 in control or 8968 +/- 3526 in AKI, %Fragments in peaks per nucleus was a mean of 59.4 +/- 10.7% in control or 56.7 +/- 12.8% in AKI. Fraction of reads in peaks, number of reads in peaks per cell and ratio of reads in genomic blacklist regions per cell for each patient were shown in Fig. S12. Differential chromatin accessibility among cell types was assessed with the Seurat FindMarkers function for peaks detected in at least 10% of cells with a likelihood ratio test and a log-fold-change threshold of 0.25. The nearby genes were determined by ClosestFeature function. Bonferroni-adjusted p-values were used to determine significance at an FDR < 0.05.

### Estimation of transcription factor activity from snATAC-seq data

Transcription factor activity was estimated using the integrated snATAC-seq dataset and chromVAR v1.10.0(*22*). The positional weight matrix was obtained from the JASPAR2022 database (collection = “CORE”, tax_group = ‘vertebrates’, all_versions = F) (*59*). Cell-type-specific chromVAR activities were calculated using the RunChromVAR wrapper in Signac v1.4.0 The chromVAR activity in each transcription factor on the whole dataset was shown with FeaturePlot function with max.cutoff = q99 and min.cutoff = q1 or VlnPlot function.

### Visualization of single nucleus dataset features

Gene expressions (snRNA-seq) or gene activities (snATAC-seq) were visualized with FeaturePlot (UMAP), VlnPlot (violin plot) or DotPlot (dot plot) function on Seurat. The fragment coverage around differentially accessible regions in each cell type was visualized by CoveragePlot function (Signac).

### Estimation of transcription factor activity from snATAC-seq data

Transcription factor activity was estimated using the integrated snATAC-seq dataset and chromVAR v1.10.0(*22*). The positional weight matrix was obtained from the JASPAR2022 database (collection = “CORE”, tax_group = ‘vertebrates’, all_versions = F) (*59*). Cell-type-specific chromVAR activities were calculated using the RunChromVAR wrapper in Signac v1.4.0 The chromVAR activity in each transcription factor on the whole dataset was shown with FeaturePlot function with max.cutoff = q99 and min.cutoff = q1 or VlnPlot function.

### Construction of pseudotemporal trajectories

Cicero v1.3.5 (*25*) was used to generate pseudotemporal trajectories for the snATAC-seq dataset. First, the cell dataset object (CDS) was constructed from the peak count matrix in the Seurat object for PT subtypes with “make_atac_cds” function with binarize = F. Next, the CDS was preprocessed (num_dim = 50), aligned to remove batch effect and reduced onto a lower dimensional space with the “reduce_dimension” function (reduction_method = ‘UMAP’, preprocess_method = “Aligned”)(*60*). After filtering potential doublets and low-quality nuclei that express non-PT cell type markers, the nuclei were clustered (cluster_cells). Subsequently, cell ordering was performed with the learn_graph function. The data were visualized with plot_cells functions.

### Construction of cis-coaccessibility network

Cicero v1.3.5 (*25*) was used to construct cis-coaccessibility network (CCAN) for the snATAC-seq FR-PTC (Fig. 4) per instructions provided on GitHub (https://cole-trapnell-lab.github.io/cicero-release/docs_m3/). Briefly, the FRPTC data was extracted from integrated snATAC-seq dataset and converted to cell dataset (CDS) objects using the make_atac_cds function. The CDS object was processed using the detect_genes() and estimate_size_factors() functions with default parameters prior to dimensional reduction and conversion to a Cicero CDS object. FRPTC-specific CCAN was generated using the run_cicero function with default parameters. CCAN was visualized with plot_connections function with coaccess_cutoff = .2 (Fig. 4).

### Cell culture

Human primary proximal tubular cells (human primary PTC, Lonza; CC-2553) were cultured with renal epithelial cell growth medium kit (Lonza; CC-3190) in a humidified 5% CO2 atmosphere at 37°C. Experiments were performed on early passages. Cells were plated at a density of 1×10^5^ cells per well in a 12-well plate, incubated overnight, and transfected with 40 pico mol siRNA for *RELA*, *RELB* or *CREB5*(ON-TARGETplus SMARTpool siRNA [Horizon Discovery], L-003533-00, L-004767-00, L-008436-00) or negative control siRNA (ON-TARGETplus Non-targeting Control Pool [Horizon Discovery], D-001810-10) using Lipofectamine RNAiMAX (Life Technologies) following the manufacturer’s protocol. Cells were harvested at 72 h after transfection for RNA isolation. For *RELA* or *RELB* knockdown, cells were also treated with or without TNFα (R&D systems, 100 ng/ml) at 48 h after transfection and harvested at 72 h after transfection (24 h after TNFα treatment).

### MTS assay

Primary human PTC were seeded at 2.5×10^3^ cells per well on 96-well tissue culture plates 24 h before transfection and transfected with 6 pico mol siRNA for *CREB5*(ON-TARGETplus SMARTpool siRNA [Horizon Discovery], L-008436-00) or negative control siRNA (ON-TARGETplus Non-targeting Control Pool [Horizon Discovery], D-001810-10) using Lipofectamine RNAiMAX (Life Technologies) following the manufacturer’s protocol. Four replicates were prepared per group. Proliferation was measured using the CellTiter96 AQueous One Solution Cell Proliferation Assay (Promega, G3582) following the manufacturer’s protocol. Optical density readings were obtained two hours following days 1, 2, 3 and 4.

### Quantitative PCR

Total RNA was extracted from primary human PTC with the Direct-zol MicroPrep Kit (Zymo) following manufacturer’s instructions. The extracted RNA (2 μg) was reverse transcribed using the High-Capacity cDNA Reverse Transcription Kit (Life Technologies). Quantitative PCR (qPCR) was performed using iTaq Universal SYBR Green Supermix (Bio-Rad). Data were normalized by the abundance of *GAPDH* mRNA. Primer sequences are shown in Table S2.

### CUT&RUN assay

We performed CUT&RUN assay with CUTANA kit (EpiCypher, 14-1048) according to the manufacturer’s instruction. The human RPTEC with early passages were seeded at 8 × 10^5^ cells on 10 cm culture dish at the day before the assay and treated with or without TNFα (R&D systems, 100 ng/ml) at 3 h before the fixation. Formaldehyde (37%, Sigma-Aldrich, 25259) was directly added to the medium of the RPTEC to achieve a final concentration of 0.5% for 1 min in room temperature. Fixation reaction was quenched by adding glycine at a final concentration of 125 mM. The cells were treated with trypsin (Gibco, 0.05%) for 2 min at 37°C, scraped from a culture dish and centrifuged at 500 × g for 5 min. The centrifugated cells were resuspended in PBS with 1% BSA and counted. 500,000 cells in 100 ul wash buffer were mixed and incubated with Concanavalin A (ConA) conjugated paramagnetic beads. Antibodies were added to each sample (RELA antibody [Santa Cruz, sc-109, 1:25] or rabbit IgG negative control antibody [EpiCypher, 13-0041, 1:50]). The remaining steps were performed according to the manufacturer’s instructions for cross-linked samples. Library preparation was performed using the NEBNext Ultra II DNA Library Prep Kit for Illumina (New England BioLabs, E7645S) with the manufacturer’s instruction, including minor modifications indicated by CUTANA kit. The libraries were sequenced on a NovaSeq instrument (Illumina, 150 bp paired-end reads). Fastq files were trimmed with Trim Galore (Cutadapt [v2.8]) and aligned with Bowtie2 [v2.3.5.1] (parameters: --local --very-sensitive-local --no-unal --no-mixed --no-discordant --phred33 -I 10 -X 700) using hg38. The SAM files were converted to BAM files with samtools (1.9). Peak calling was performed using MACS2 [v2.2.7.1] with default parameters. Visualization was performed with Integrated Genome Viewer(*61*) and bigWig files generated from BAM files using DeepTools (3.5.0). The visualization for modified histone (H3K4me3 and H3K27ac) in Fig. 3A was generated with previously published CUT&RUN data for human primary PTC(*44*). CUT&RUN peaks were annotated with ChIPSeeker (v1.24.0)(*62*) using hg38. To convert the genome coordinates for human RELA binding sites (hg38) to thosee of mouse genome (mm10), The Genome Browser Convert utility (https://genome.ucsc.edu/cgi-bin/hgLiftOver) was used with Minimum ratio of bases that must remap = 0.1 (*31*).

### Reanalysis of bulk RNA-seq

The bulk RNA-seq dataset for primary human PTC with siRNA knockdown of *CREB5* or control was retrieved from GEO220222(*44*). Reads were then aligned with STAR v2.7.9a to the Ensembl release 101 primary assembly. Gene counts were calculated from the number of uniquely aligned, unambiguous reads by Subread:featureCount v2.0.3.

### Gene set enrichment analysis and gene ontology analysis

Single nucleus gene set enrichment analysis was performed with the VISION v2.1.0 R package according to instructions provided on GitHub (https://github.com/YosefLab/VISION)(43), using Hallmark gene sets obtained from the Molecular Signatures Database v7.4 distributed at the GSEA Web site. The heatmaps were generated with pheatmap v1.0.12 from gene set enrichment scores averaged in each cell type (Fig. 6C). The bulk RNA-seq data was analyzed with GSEA v4.0.3 software (Broad Institute) (*63*) with Hallmark gene sets. The top-ranked gene set list in Fig. 7E only displays those with false discovery rate (FDR) < 0.05. Gene ontology (GO) term analysis was performed with Gene Ontology Resource (https://geneontology.org/) (*64*, *65*).

### Gene regulatory network analysis with RENIN

To construct gene regulatory networks, we first generated cis-coaccessibility networks (CCAN) in our snATAC-seq datasets with Cicero v1.3.9(*25*). Within each CCAN, we identified peaks that overlapped with the 2 kbp promoter region or gene body for genes that were differentially expressed between the PT and iPT clusters in each of control and AKI group in the human snRNA-seq dataset or between the healthy PT (PTS1-3) and acutely injured PT (New PT1) or FR-PTC (New PT2) clusters in the mouse snRNA-seq dataset (*8*). Differentially expressed genes with linked CCANs were included for TF modeling. For each species, we then used the filtered cisBP v0.2 motif database provided by the chromVARmotifs package and JASPAR2022 to identify predicted transcription factor (TF) binding sites within each differentially expressed gene-linked CCAN. A preliminary list of TFs that regulate each differentially expressed gene was generated by aggregating TFs with at least one predicted binding motif within each gene’s CCAN. From the preliminary list of TFs regulating each differentially expressed gene, we then used the adaptive elastic net-based RENIN to generate parametric gene regulatory networks for each gene. We then ranked TFs by the sum of their coefficients across differentially expressed genes between groups, multiplied by each TF’s mean proximal tubule expression. Regulatory coefficients for differentially expressed genes that were upregulated in the normal PT cluster versus the injury-associated cluster were multiplied by -1 to allow for identification of failed repair or injury association for each TF. To evaluate interspecies RENIN scores (Fig. 7A), the mouse genes were converted to human genes using biomaRt and ensembl with getLDS function.

### Statistical analysis

No statistical methods were used to predetermine sample size for single nucleus analysis. Experiments were not randomized and investigators were not blinded to allocation during library preparation, experiments or analysis. Bonferroni adjusted *P* values were used to determine significance for differential gene expression or accessibility. The quantitative PCR data (Fig. 4, 7 and Fig. S8) are presented as mean±s.d. and were compared between groups with two-sided Student’s *t*-test (Fig. 7) or one-way ANOVA with post hoc Turkey test (Fig. 4, Fig. S8). The MTS assay data were compared between groups with two-sided Student’s *t*-test at each time point (Fig.7). A *P* value of <0.05 was considered statistically significant.

## Funding

These experiments were funded by NIDDK UC2DK126024, 2R01DK103740 and 1U54DK137332 (B.D.H.). Additional support was from the Japan Society for the Promotion of Science (JSPS) Postdoctoral Fellowships for Research Abroad, The Osamu Hayaishi Memorial Scholarship for Study Abroad and American Society of Nephrology Carl W. Gottschalk Research Scholar Award (Y.M.)

## Author contributions

Conceptualization: BDH

Library preparation: EED, YY

Computational analysis: YM, NL, HW

Experiment: YM, EED, YY, NL, YK

Online tool: HW

Supervision: BDH Writing: YM, BDH

## Competing interests

B.D.H. is a consultant for Janssen Research & Development, LLC, Pfizer and Chinook Therapeutics, holds equity in Chinook Therapeutics and grant funding from Chinook Therapeutics and Janssen Research & Development. Y.Y. is currently an employee of Chugai Pharmaceutical Co., Ltd. The remaining authors declare no competing interests.

## Notes

### Summary of Updates

Minor fixations for authorship-related matter.

## References

1. A. S. Levey, M. T. James, Acute Kidney Injury. Ann Intern Med 167, ITC66–ITC80 (2017).

2. C. Ronco, R. Bellomo, J. A. Kellum, Acute kidney injury. Lancet 394, 1949–1964 (2019).

3. L. S. Chawla, P. W. Eggers, R. A. Star, P. L. Kimmel, Acute kidney injury and chronic kidney disease as interconnected syndromes. N Engl J Med 371, 58–66 (2014).

4. T. Chen, Q. Cao, Y. Wang, D. C. H. Harris, M2 macrophages in kidney disease: biology, therapies, and perspectives. Kidney Int 95, 760–773 (2019).

5. H. I. Han, L. B. Skvarca, E. B. Espiritu, A. J. Davidson, N. A. Hukriede, The role of macrophages during acute kidney injury: destruction and repair. Pediatr Nephrol 34, 561– 569 (2019).

6. B. D. Humphreys, Mechanisms of Renal Fibrosis. Annu Rev Physiol 80, 309–326 (2018).

7. E. R. Gibney, C. M. Nolan, Epigenetics and gene expression. Heredity (Edinb) 105, 4–13 (2010).

8. Y. Kirita, H. Wu, K. Uchimura, P. C. Wilson, B. D. Humphreys, Cell profiling of mouse acute kidney injury reveals conserved cellular responses to injury. Proc Natl Acad Sci U S A 117, 15874–15883 (2020).

9. L. M. S. Gerhardt, J. Liu, K. Koppitch, P. E. Cippà, A. P. McMahon, Single-nuclear transcriptomics reveals diversity of proximal tubule cell states in a dynamic response to acute kidney injury. Proc Natl Acad Sci U S A 118, e2026684118 (2021).

10. S. Ide, Y. Kobayashi, K. Ide, S. A. Strausser, K. Abe, S. Herbek, L. L. O’Brien, S. D. Crowley, L. Barisoni, A. Tata, P. R. Tata, T. Souma, Ferroptotic stress promotes the accumulation of pro-inflammatory proximal tubular cells in maladaptive renal repair. Elife 10, e68603 (2021).

11. Y. Muto, P. C. Wilson, N. Ledru, H. Wu, H. Dimke, S. S. Waikar, B. D. Humphreys, Single cell transcriptional and chromatin accessibility profiling redefine cellular heterogeneity in the adult human kidney. Nat Commun 12, 2190 (2021).

12. Y. Muto, E. E. Dixon, Y. Yoshimura, H. Wu, K. Omachi, N. Ledru, P. C. Wilson, A. J. King, N. Eric Olson, M. G. Gunawan, J. J. Kuo, J. H. Cox, J. H. Miner, S. L. Seliger, O. M. Woodward, P. A. Welling, T. J. Watnick, B. D. Humphreys, Defining cellular complexity in human autosomal dominant polycystic kidney disease by multimodal single cell analysis. Nat Commun 13, 6497 (2022).

13. P. C. Wilson, Y. Muto, H. Wu, A. Karihaloo, S. S. Waikar, B. D. Humphreys, Multimodal single cell sequencing implicates chromatin accessibility and genetic background in diabetic kidney disease progression. Nat Commun 13, 5253 (2022).

14. Y.-H. Chou, S.-Y. Pan, Y.-H. Shao, H.-M. Shih, S.-Y. Wei, C.-F. Lai, W.-C. Chiang, C. Schrimpf, K.-C. Yang, L.-C. Lai, Y.-M. Chen, T.-S. Chu, S.-L. Lin, Methylation in pericytes after acute injury promotes chronic kidney disease. J Clin Invest 130, 4845–4857 (2020).

15. M. Nangaku, Y. Hirakawa, I. Mimura, R. Inagi, T. Tanaka, Epigenetic Changes in the Acute Kidney Injury-to-Chronic Kidney Disease Transition. Nephron 137, 256–259 (2017).

16. X. Cao, J. Wang, T. Zhang, Z. Liu, L. Liu, Y. Chen, Z. Li, Y. Zhao, Q. Yu, T. Liu, J. Nie, Y. Niu, Y. Chen, L. Yang, L. Zhang, Chromatin accessibility dynamics dictate renal tubular epithelial cell response to injury. Nat Commun 13, 7322 (2022).

17. L. M. S. Gerhardt, K. Koppitch, J. van Gestel, J. Guo, S. Cho, H. Wu, Y. Kirita, B. D. Humphreys, A. P. McMahon, Lineage Tracing and Single-Nucleus Multiomics Reveal Novel Features of Adaptive and Maladaptive Repair after Acute Kidney Injury. J Am Soc Nephrol 34, 554–571 (2023).

18. Y. Hao, S. Hao, E. Andersen-Nissen, W. M. Mauck, S. Zheng, A. Butler, M. J. Lee, A. J. Wilk, C. Darby, M. Zager, P. Hoffman, M. Stoeckius, E. Papalexi, E. P. Mimitou, J. Jain, A. Srivastava, T. Stuart, L. M. Fleming, B. Yeung, A. J. Rogers, J. M. McElrath, C. A. Blish, R. Gottardo, P. Smibert, R. Satija, Integrated analysis of multimodal single-cell data. Cell 184, 3573–3587.e29 (2021).

19. T. Stuart, A. Srivastava, S. Madad, C. A. Lareau, R. Satija, Single-cell chromatin state analysis with Signac. Nat Methods 18, 1333–1341 (2021).

20. I. Korsunsky, N. Millard, J. Fan, K. Slowikowski, F. Zhang, K. Wei, Y. Baglaenko, M. Brenner, P. Loh, S. Raychaudhuri, Fast, sensitive and accurate integration of single-cell data with Harmony. Nature Methods 16, 1289–1296 (2019).

21. J. Liu, S. Kumar, E. Dolzhenko, G. F. Alvarado, J. Guo, C. Lu, Y. Chen, M. Li, M. C. Dessing, R. K. Parvez, P. E. Cippà, A. M. Krautzberger, G. Saribekyan, A. D. Smith, A. P. McMahon, Molecular characterization of the transition from acute to chronic kidney injury following ischemia/reperfusion. JCI Insight 2 (2017).

22. A. N. Schep, B. Wu, J. D. Buenrostro, W. J. Greenleaf, chromVAR: inferring transcription-factor-associated accessibility from single-cell epigenomic data. Nat Methods 14, 975–978 (2017).

23. L. M. Shelton, B. K. Park, I. M. Copple, Role of Nrf2 in protection against acute kidney injury. Kidney Int 84, 1090–1095 (2013).

24. C.-T. Ong, V. G. Corces, CTCF: an architectural protein bridging genome topology and function. Nat Rev Genet 15, 234–246 (2014).

25. H. A. Pliner, J. S. Packer, J. L. McFaline-Figueroa, D. A. Cusanovich, R. M. Daza, D. Aghamirzaie, S. Srivatsan, X. Qiu, D. Jackson, A. Minkina, A. C. Adey, F. J. Steemers, J. Shendure, C. Trapnell, Cicero Predicts cis-Regulatory DNA Interactions from Single-Cell Chromatin Accessibility Data. Mol. Cell 71, 858–871.e8 (2018).

26. M. Bleu, S. Gaulis, R. Lopes, K. Sprouffske, V. Apfel, S. Holwerda, M. Pregnolato, U. Yildiz, V. Cordoʹ, A. F. M. Dost, J. Knehr, W. Carbone, F. Lohmann, C. Y. Lin, J. E. Bradner, A. Kauffmann, L. Tordella, G. Roma, G. G. Galli, PAX8 activates metabolic genes via enhancer elements in Renal Cell Carcinoma. Nat Commun 10, 3739 (2019).

27. A. S. A. Patel, S. Hirosue, P. Rodrigues, E. Vojtasova, E. K. Richardson, J. Ge, S. E. Syafruddin, Speed, E. K. Papachristou, D. Baker, D. Clarke, S. Purvis, L. Wesolowski, A. Dyas, L. Castillon, V. Caraffini, D. Bihary, C. Yong, D. J. Harrison, G. D. Stewart, M. J. Machiela, M. P. Purdue, S. J. Chanock, A. Y. Warren, S. A. Samarajiwa, J. S. Carroll, S. Vanharanta, The renal lineage factor PAX8 controls oncogenic signalling in kidney cancer. Nature 606, 999–1006 (2022).

28. A. J. Peired, G. Antonelli, M. L. Angelotti, M. Allinovi, F. Guzzi, A. Sisti, R. Semeraro, C. Conte, B. Mazzinghi, S. Nardi, M. E. Melica, L. De Chiara, E. Lazzeri, L. Lasagni, T. Lottini, S. Landini, S. Giglio, A. Mari, F. Di Maida, A. Antonelli, F. Porpiglia, R. Schiavina, V. Ficarra, D. Facchiano, M. Gacci, S. Serni, M. Carini, G. J. Netto, R. M. Roperto, A. Magi, C. F. Christiansen, M. Rotondi, H. Liapis, H.-J. Anders, A. Minervini, M. R. Raspollini, P. Romagnani, Acute kidney injury promotes development of papillary renal cell adenoma and carcinoma from renal progenitor cells. Sci Transl Med 12, eaaw6003 (2020).

29. P. J. Skene, S. Henikoff, An efficient targeted nuclease strategy for high-resolution mapping of DNA binding sites. Elife 6, e21856 (2017).

30. M. S. Hayden, S. Ghosh, Regulation of NF-κB by TNF family cytokines. Semin Immunol 26, 253–266 (2014).

31. W. J. Kent, C. W. Sugnet, T. S. Furey, K. M. Roskin, T. H. Pringle, A. M. Zahler, D. Haussler, The human genome browser at UCSC. Genome Res 12, 996–1006 (2002).

32. L. Xu, D. Sharkey, L. G. Cantley, Tubular GM-CSF Promotes Late MCP-1/CCR2-Mediated Fibrosis and Inflammation after Ischemia/Reperfusion Injury. J Am Soc Nephrol 30, 1825– 1840 (2019).

33. L. Li, L. Huang, S.-S. J. Sung, A. L. Vergis, D. L. Rosin, C. E. Rose, P. I. Lobo, M. D. Okusa, The chemokine receptors CCR2 and CX3CR1 mediate monocyte/macrophage trafficking in kidney ischemia-reperfusion injury. Kidney Int 74, 1526–1537 (2008).

34. I. Ushach, A. Zlotnik, Biological role of granulocyte macrophage colony-stimulating factor (GM-CSF) and macrophage colony-stimulating factor (M-CSF) on cells of the myeloid lineage. J Leukoc Biol 100, 481–489 (2016).

35. M.-Z. Zhang, B. Yao, S. Yang, L. Jiang, S. Wang, X. Fan, H. Yin, K. Wong, T. Miyazawa, J. Chen, I. Chang, A. Singh, R. C. Harris, CSF-1 signaling mediates recovery from acute kidney injury. J Clin Invest 122, 4519–4532 (2012).

36. Y. Wang, J. Chang, B. Yao, A. Niu, E. Kelly, M. C. Breeggemann, S. L. Abboud Werner, R. C. Harris, M.-Z. Zhang, Proximal tubule-derived colony stimulating factor-1 mediates polarization of renal macrophages and dendritic cells, and recovery in acute kidney injury. Kidney Int 88, 1274–1282 (2015).

37. J. Menke, Y. Iwata, W. A. Rabacal, R. Basu, Y. G. Yeung, B. D. Humphreys, T. Wada, A. Schwarting, E. R. Stanley, V. R. Kelley, CSF-1 signals directly to renal tubular epithelial cells to mediate repair in mice. J Clin Invest 119, 2330–2342 (2009).

38. L. Meziani, M. Mondini, B. Petit, A. Boissonnas, V. Thomas de Montpreville, O. Mercier, M.-C. Vozenin, E. Deutsch, CSF1R inhibition prevents radiation pulmonary fibrosis by depletion of interstitial macrophages. Eur Respir J 51, 1702120 (2018).

39. N. Joshi, S. Watanabe, R. Verma, R. P. Jablonski, C.-I. Chen, P. Cheresh, N. S. Markov, P. A. Reyfman, A. C. McQuattie-Pimentel, L. Sichizya, Z. Lu, R. Piseaux-Aillon, D. Kirchenbuechler, A. S. Flozak, C. J. Gottardi, C. M. Cuda, H. Perlman, M. Jain, D. W. Kamp, G. R. S. Budinger, A. V. Misharin, A spatially restricted fibrotic niche in pulmonary fibrosis is sustained by M-CSF/M-CSFR signalling in monocyte-derived alveolar macrophages. Eur Respir J 55, 1900646 (2020).

40. S. Jaiswal, M. P. Chao, R. Majeti, I. L. Weissman, Macrophages as mediators of tumor immunosurveillance. Trends Immunol 31, 212–219 (2010).

41. J. Barrera-Chimal, G. R. Estrela, S. M. Lechner, S. Giraud, S. El Moghrabi, S. Kaaki, P. Kolkhof, T. Hauet, F. Jaisser, The myeloid mineralocorticoid receptor controls inflammatory and fibrotic responses after renal injury via macrophage interleukin-4 receptor signaling. Kidney Int 93, 1344–1355 (2018).

42. J. M. Luther, A. B. Fogo, The role of mineralocorticoid receptor activation in kidney inflammation and fibrosis. Kidney International Supplements 12, 63–68 (2022).

43. D. DeTomaso, M. G. Jones, M. Subramaniam, T. Ashuach, C. J. Ye, N. Yosef, Functional interpretation of single cell similarity maps. Nat Commun 10, 4376 (2019).

44. N. Ledru, P. C. Wilson, Y. Muto, Y. Yoshimura, H. Wu, A. Asthana, S. G. Tullius, S. S. Waikar, G. Orlando, B. D. Humphreys, “Predicting regulators of epithelial cell state through regularized regression analysis of single cell multiomic sequencing” (preprint, Genomics, 2022); 10.1101/2022.12.29.522232.

45. S. E. Piret, Y. Guo, A. A. Attallah, S. J. Horne, A. Zollman, D. Owusu, J. Henein, V. S. Sidorenko, M. P. Revelo, T. Hato, A. Ma’ayan, J. C. He, S. K. Mallipattu, Krüppel-like factor 6-mediated loss of BCAA catabolism contributes to kidney injury in mice and humans. Proc Natl Acad Sci U S A 118, e2024414118 (2021).

46. H. S. Kang, J. Y. Beak, Y.-S. Kim, R. Herbert, A. M. Jetten, Glis3 is associated with primary cilia and Wwtr1/TAZ and implicated in polycystic kidney disease. Mol Cell Biol 29, 2556– 2569 (2009).

47. V. Senée, C. Chelala, S. Duchatelet, D. Feng, H. Blanc, J.-C. Cossec, C. Charon, M. Nicolino, P. Boileau, D. R. Cavener, P. Bougnères, D. Taha, C. Julier, Mutations in GLIS3 are responsible for a rare syndrome with neonatal diabetes mellitus and congenital hypothyroidism. Nat Genet 38, 682–687 (2006).

48. S. Wang, J. Qiu, L. Liu, C. Su, L. Qi, C. Huang, X. Chen, Y. Zhang, Y. Ye, Y. Ding, L. Liang, W. Liao, CREB5 promotes invasiveness and metastasis in colorectal cancer by directly activating MET. J Exp Clin Cancer Res 39, 168 (2020).

49. S. He, Y. Deng, Y. Liao, X. Li, J. Liu, S. Yao, CREB5 promotes tumor cell invasion and correlates with poor prognosis in epithelial ovarian cancer. Oncol Lett 14, 8156–8161 (2017).

50. J. Wu, S.-T. Wang, Z.-J. Zhang, Q. Zhou, B.-G. Peng, CREB5 promotes cell proliferation and correlates with poor prognosis in hepatocellular carcinoma. Int J Clin Exp Pathol 11, 4908– 4916 (2018).

51. J. H. Hwang, J.-H. Seo, M. L. Beshiri, S. Wankowicz, D. Liu, A. Cheung, J. Li, X. Qiu, A. L. Hong, G. Botta, L. Golumb, C. Richter, J. So, G. J. Sandoval, A. O. Giacomelli, S. H. Ly, C. Han, C. Dai, H. Pakula, A. Sheahan, F. Piccioni, O. Gjoerup, M. Loda, A. G. Sowalsky, L. Ellis, H. Long, D. E. Root, K. Kelly, E. M. Van Allen, M. L. Freedman, A. D. Choudhury, W. C. Hahn, CREB5 Promotes Resistance to Androgen-Receptor Antagonists and Androgen Deprivation in Prostate Cancer. Cell Rep 29, 2355–2370.e6 (2019).

52. W. Shi, W. Le, Q. Tang, S. Shi, J. Shi, Regulon analysis identifies protective FXR and CREB5 in proximal tubules in early diabetic kidney disease. BMC Nephrol 24, 180 (2023).

53. L. Markó, E. Vigolo, C. Hinze, J.-K. Park, G. Roël, A. Balogh, M. Choi, A. Wübken, J. Cording, I. E. Blasig, F. C. Luft, C. Scheidereit, K. M. Schmidt-Ott, R. Schmidt-Ullrich, D. N. Müller, Tubular Epithelial NF-κB Activity Regulates Ischemic AKI. JASN 27, 2658– 2669 (2016).

54. L. Yang, C. R. Brooks, S. Xiao, V. Sabbisetti, M. Y. Yeung, L.-L. Hsiao, T. Ichimura, V. Kuchroo, J. V. Bonventre, KIM-1-mediated phagocytosis reduces acute injury to the kidney. J Clin Invest 125, 1620–1636 (2015).

55. T. Stuart, A. Srivastava, C. Lareau, R. Satija, “Multimodal single-cell chromatin analysis with Signac” (preprint, Genomics, 2020); 10.1101/2020.11.09.373613.

56. A. Thibodeau, A. Eroglu, C. S. McGinnis, N. Lawlor, D. Nehar-Belaid, R. Kursawe, R. Marches, D. N. Conrad, G. A. Kuchel, Z. J. Gartner, J. Banchereau, M. L. Stitzel, A. E. Cicek, D. Ucar, AMULET: a novel read count-based method for effective multiplet detection from single nucleus ATAC-seq data. Genome Biol 22, 252 (2021).

57. M. D. Young, S. Behjati, SoupX removes ambient RNA contamination from droplet-based single-cell RNA sequencing data. Gigascience 9, giaa151 (2020).

58. C. S. McGinnis, L. M. Murrow, Z. J. Gartner, DoubletFinder: Doublet Detection in Single-Cell RNA Sequencing Data Using Artificial Nearest Neighbors. Cell Systems 8, 329–337.e4 (2019).

59. J. A. Castro-Mondragon, R. Riudavets-Puig, I. Rauluseviciute, R. Berhanu Lemma, L. Turchi, R. Blanc-Mathieu, J. Lucas, P. Boddie, A. Khan, N. Manosalva Pérez, O. Fornes, T. Y. Leung, A. Aguirre, F. Hammal, D. Schmelter, D. Baranasic, B. Ballester, A. Sandelin, B. Lenhard, K. Vandepoele, W. W. Wasserman, F. Parcy, A. Mathelier, JASPAR 2022: the 9th release of the open-access database of transcription factor binding profiles. Nucleic Acids Research 50, D165–D173 (2022).

60. L. Haghverdi, A. T. L. Lun, M. D. Morgan, J. C. Marioni, Batch effects in single-cell RNA-sequencing data are corrected by matching mutual nearest neighbors. Nat Biotechnol 36, 421–427 (2018).

61. J. T. Robinson, H. Thorvaldsdóttir, W. Winckler, M. Guttman, E. S. Lander, G. Getz, J. P. Mesirov, Integrative genomics viewer. Nat Biotechnol 29, 24–26 (2011).

62. G. Yu, L.-G. Wang, Q.-Y. He, ChIPseeker: an R/Bioconductor package for ChIP peak annotation, comparison and visualization. Bioinformatics 31, 2382–2383 (2015).

63. A. Subramanian, P. Tamayo, V. K. Mootha, S. Mukherjee, B. L. Ebert, M. A. Gillette, A. Paulovich, S. L. Pomeroy, T. R. Golub, E. S. Lander, J. P. Mesirov, Gene set enrichment analysis: A knowledge-based approach for interpreting genome-wide expression profiles. Proceedings of the National Academy of Sciences 102, 15545–15550 (2005).

64. M. Ashburner, C. A. Ball, J. A. Blake, D. Botstein, H. Butler, J. M. Cherry, A. P. Davis, K. Dolinski, S. S. Dwight, J. T. Eppig, M. A. Harris, D. P. Hill, L. Issel-Tarver, A. Kasarskis, S. Lewis, J. C. Matese, J. E. Richardson, M. Ringwald, G. M. Rubin, G. Sherlock, Gene Ontology: tool for the unification of biology. Nat Genet 25, 25–29 (2000).

65. Gene Ontology Consortium, S. A. Aleksander, J. Balhoff, S. Carbon, J. M. Cherry, H. J. Drabkin, D. Ebert, M. Feuermann, P. Gaudet, N. L. Harris, D. P. Hill, R. Lee, H. Mi, S. Moxon, C. J. Mungall, A. Muruganugan, T. Mushayahama, P. W. Sternberg, P. D. Thomas, K. Van Auken, J. Ramsey, D. A. Siegele, R. L. Chisholm, P. Fey, M. C. Aspromonte, M. V. Nugnes, F. Quaglia, S. Tosatto, M. Giglio, S. Nadendla, G. Antonazzo, H. Attrill, G. Dos Santos, S. Marygold, V. Strelets, C. J. Tabone, J. Thurmond, P. Zhou, S. H. Ahmed, P. Asanitthong, D. Luna Buitrago, M. N. Erdol, M. C. Gage, M. Ali Kadhum, K. Y. C. Li, M. Long, A. Michalak, A. Pesala, A. Pritazahra, S. C. C. Saverimuttu, R. Su, K. E. Thurlow, R. C. Lovering, C. Logie, S. Oliferenko, J. Blake, K. Christie, L. Corbani, M. E. Dolan, H. J. Drabkin, D. P. Hill, L. Ni, D. Sitnikov, C. Smith, A. Cuzick, J. Seager, L. Cooper, J. Elser, P. Jaiswal, P. Gupta, P. Jaiswal, S. Naithani, M. Lera-Ramirez, K. Rutherford, V. Wood, J. L. De Pons, M. R. Dwinell, G. T. Hayman, M. L. Kaldunski, A. E. Kwitek, S. J. F. Laulederkind, M. A. Tutaj, M. Vedi, S.-J. Wang, P. D’Eustachio, L. Aimo, K. Axelsen, A. Bridge, N. Hyka-Nouspikel, A. Morgat, S. A. Aleksander, J. M. Cherry, S. R. Engel, K. Karra, S. R. Miyasato, R. S. Nash, M. S. Skrzypek, S. Weng, E. D. Wong, E. Bakker, T. Z. Berardini, L. Reiser, A. Auchincloss, K. Axelsen, G. Argoud-Puy, M.-C. Blatter, E. Boutet, L. Breuza, A. Bridge, C. Casals-Casas, E. Coudert, A. Estreicher, M. Livia Famiglietti, M. Feuermann, A. Gos, N. Gruaz-Gumowski, C. Hulo, N. Hyka-Nouspikel, F. Jungo, P. Le Mercier, D. Lieberherr, P. Masson, A. Morgat, I. Pedruzzi, L. Pourcel, S. Poux, C. Rivoire, S. Sundaram, A. Bateman, E. Bowler-Barnett, H. Bye-A-Jee, P. Denny, A. Ignatchenko, R. Ishtiaq, A. Lock, Y. Lussi, M. Magrane, M. J. Martin, S. Orchard, P. Raposo, E. Speretta, N. Tyagi, K. Warner, R. Zaru, A. D. Diehl, R. Lee, J. Chan, S. Diamantakis, D. Raciti, M. Zarowiecki, M. Fisher, C. James-Zorn, V. Ponferrada, A. Zorn, S. Ramachandran, L. Ruzicka, M. Westerfield, The Gene Ontology knowledgebase in 2023. Genetics 224, iyad031 (2023).

