## Supplemental Materials for "Epigenetic reprogramming driving successful and failed repair in acute kidney injury"

Yoshiharu Muto *et al.*

**This PDF file includes:**

Figs. S1 to S17
Tables S1 to S2

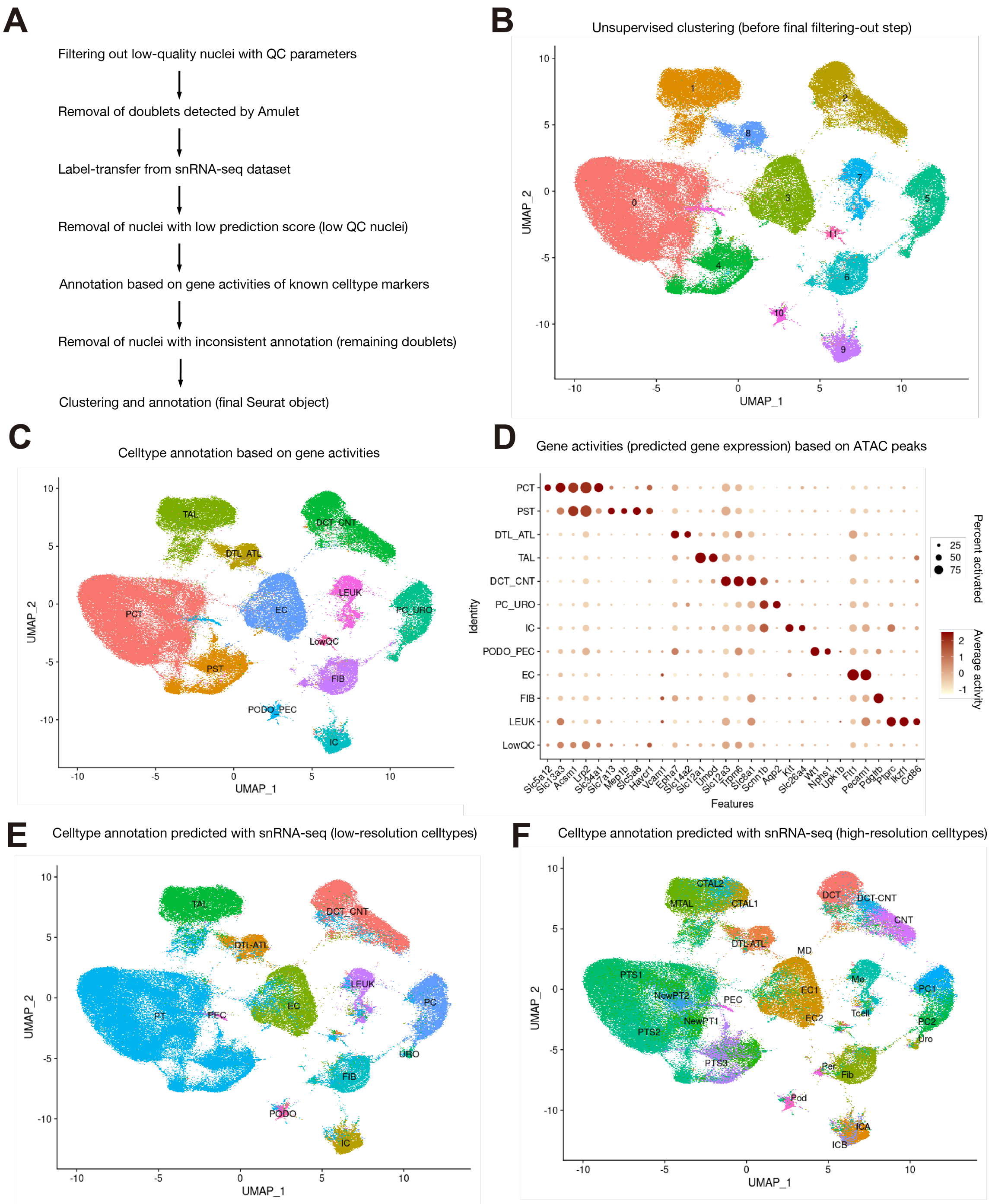


**Fig. S1. Preprocessing strategy for mouse kidney snATAC-seq atlas**
(**A**) Overview of preprocessing strategy. (**B**) UMAP plot for aggregated snATAC-seq dataset before the final filtering-out step to remove the remaining doublets and low-QC nuclei. (**C**) UMAP plot with cell type annotations based on gene activities. (**D**) Dot plot showing gene activities of cell -type marker genes for (C). (**E, F**) UMAP plot with cell type annotation predictions using snRNA-seq dataset (Fig. 1B, GSE139107) with low-resolution cell types (E) or high-resolution cell types (F). The nuclei with inconsistent annotations between gene activity-based (B) and low-resolution cell type prediction (E) were removed as remaining doublets (A). See also Method.

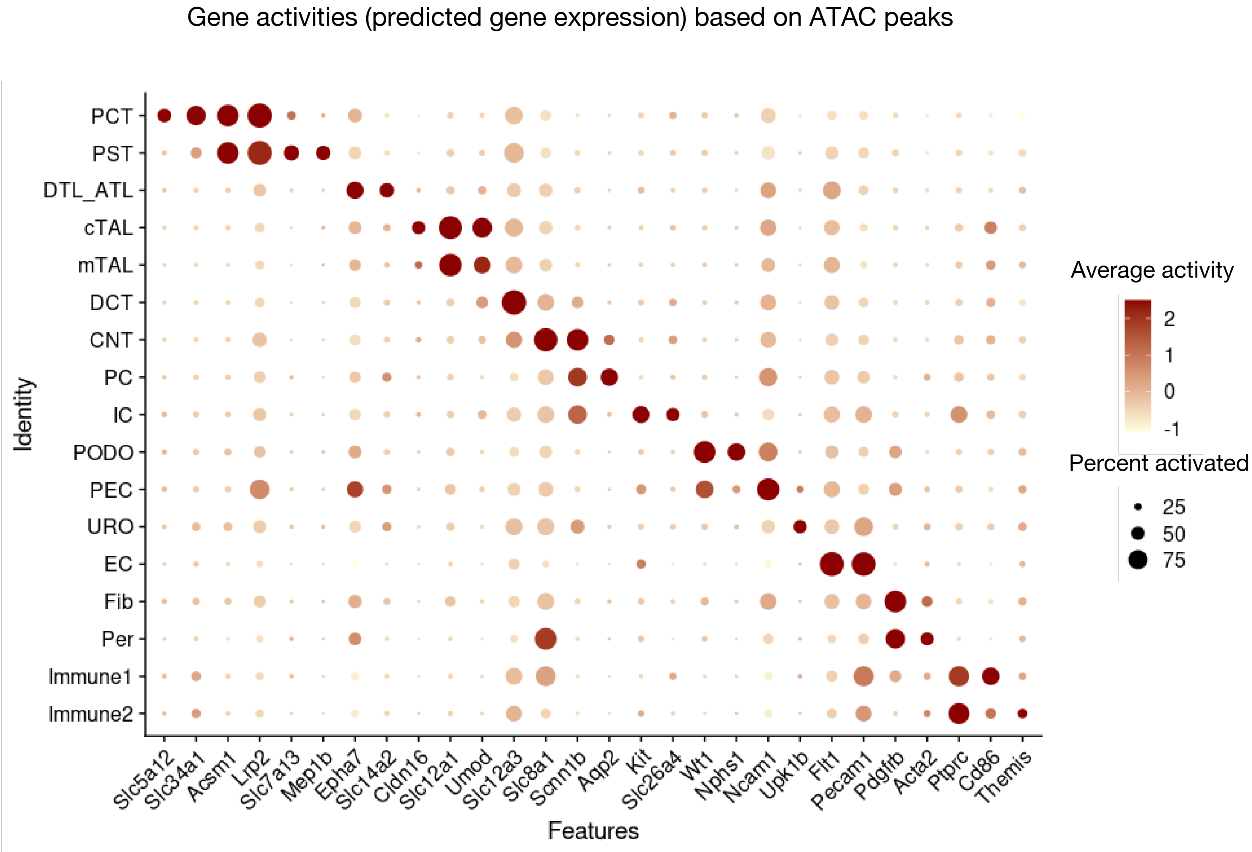


**Fig. S2. Gene activities for mouse kidney snATAC-seq atlas**
Dot plot showing gene activities calculated with snATAC-seq peaks on the gene promoter and gene body. The diameter of the dot corresponds to the proportion of nuclei with detected activity of indicated gene and the density of the dot corresponds to average gene activity relative to all cell types.

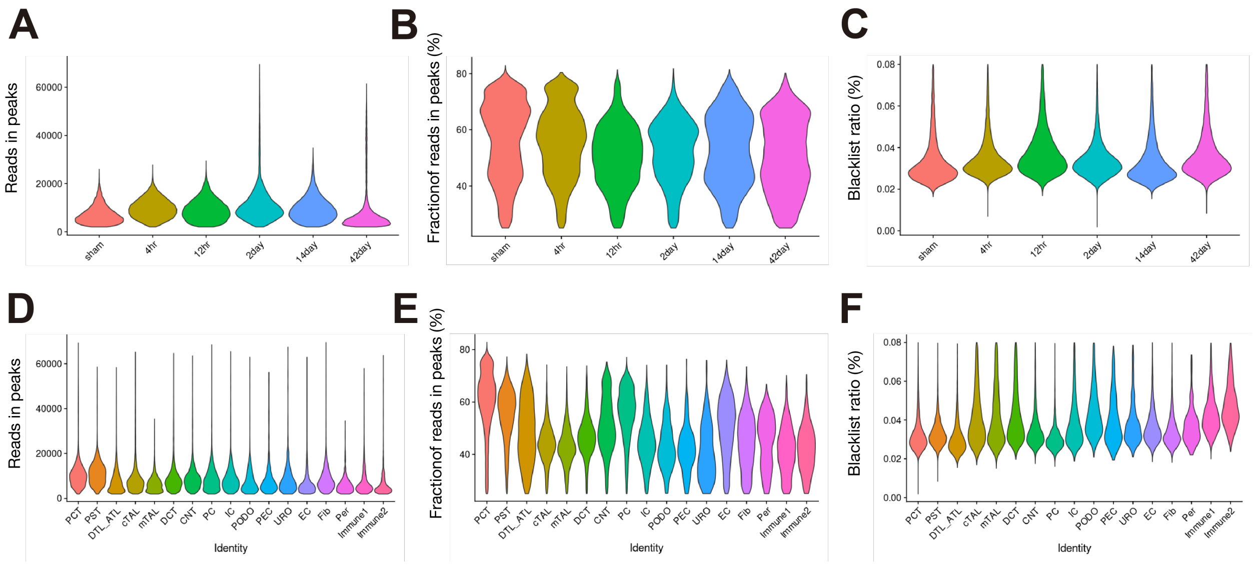


**Fig. S3. Quality metrics for mouse kidney snATAC-seq atlas**

(**A**-**C**) Violin plots showing: (**A**) number of reads in peaks per nucleus, (**B**) fraction of reads in peaks and (**C**) ratio of reads in genomic blacklist region per nucleus in each time point of snATAC-seq data. (**D**-**F**) Violin plots showing: (**D**) number of reads in peaks per nucleus, (**E**) fraction of reads in peaks and (**F**) ratio of reads in genomic blacklist region per nucleus in each cell type.

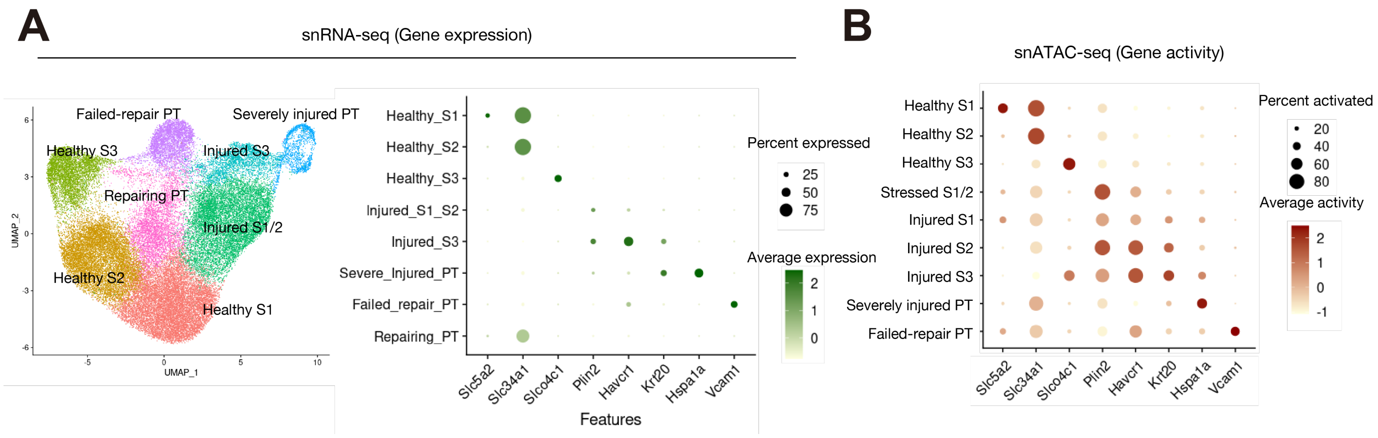


**Fig. S4. Gene expressions and activities in proximal tubular cell subtypes**

(**A**) UMAP plot (left) and dot plot showing gene expression pattern (right) for mouse proximal tubular cells in previously published snRNA-seq dataset (GSE139107) for mouse kidneys with IRI. Subclustering was performed in our previous study by Kirita et al (*8*). (**B**) Dot plot showing gene activity pattern for subclustering of PT cells in snATAC-seq (Fig. 2A) for mouse kidneys with IRI.

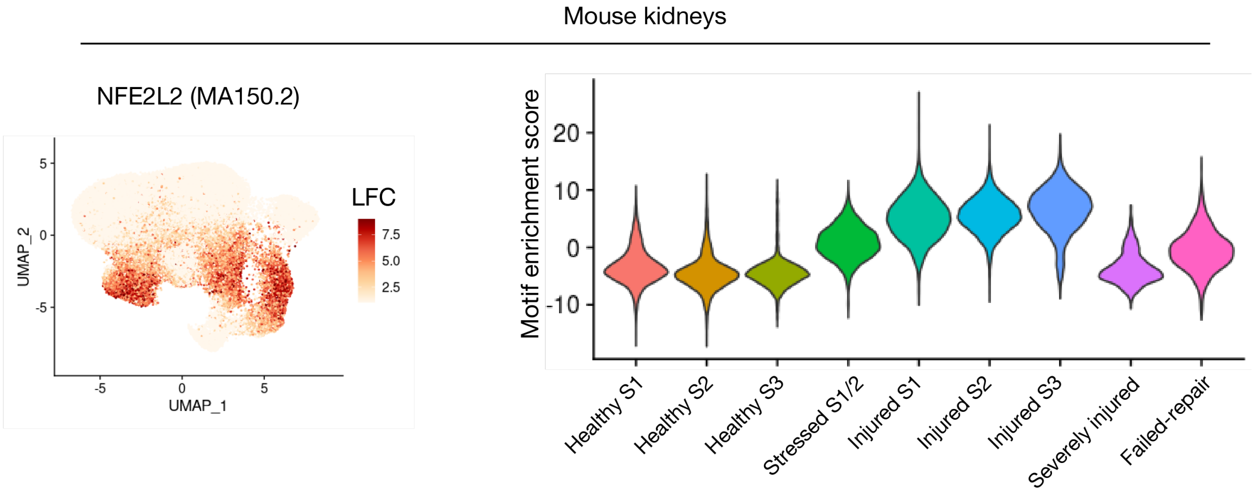


**Fig. S5. NFE2L2 motif enrichment among proximal tubular cell subtypes**
UMAP plot (left) and violin plot (right) displaying transcription factor motif enrichment among PT subtypes for NFE2L2 (MA0150.2) in mouse kidneys. The color scale for the UMAP plot represents a normalized log-fold-change (LFC).

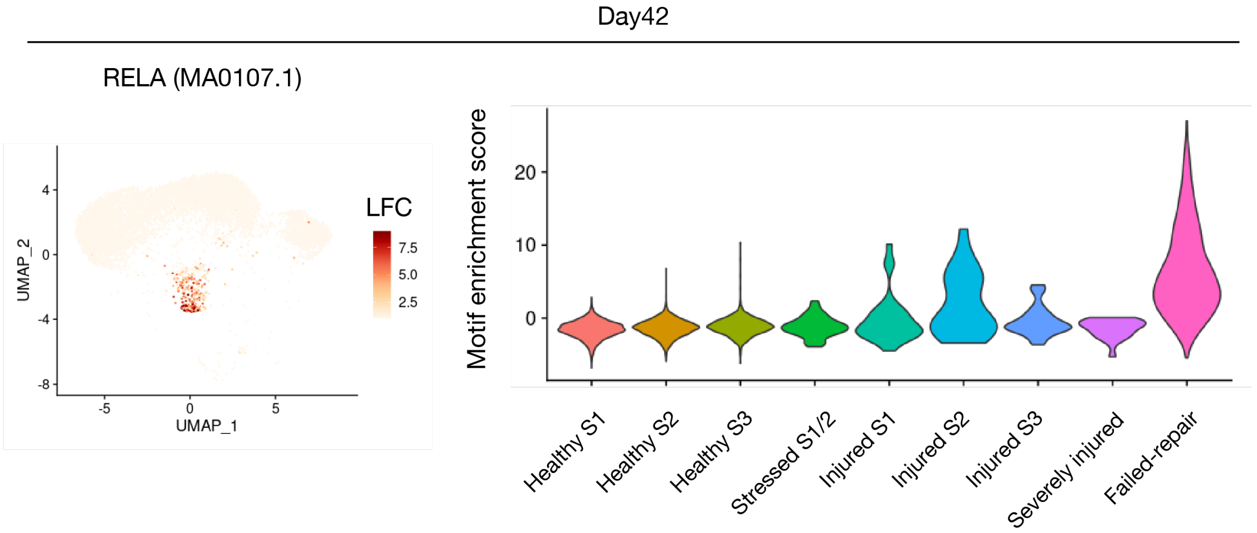


**Fig. S6. RELA motif enrichment among proximal tubular cell subtypes at Day42**
UMAP plot (left) and violin plot (right) displaying transcription factor motif enrichment among PT subtypes for RELA (MA0107.1) in mouse kidneys at day 42 following IRI. RELA motif was most enriched among failed-repair PT cells. The color scale for the UMAP plot represents a normalized log-fold-change (LFC).

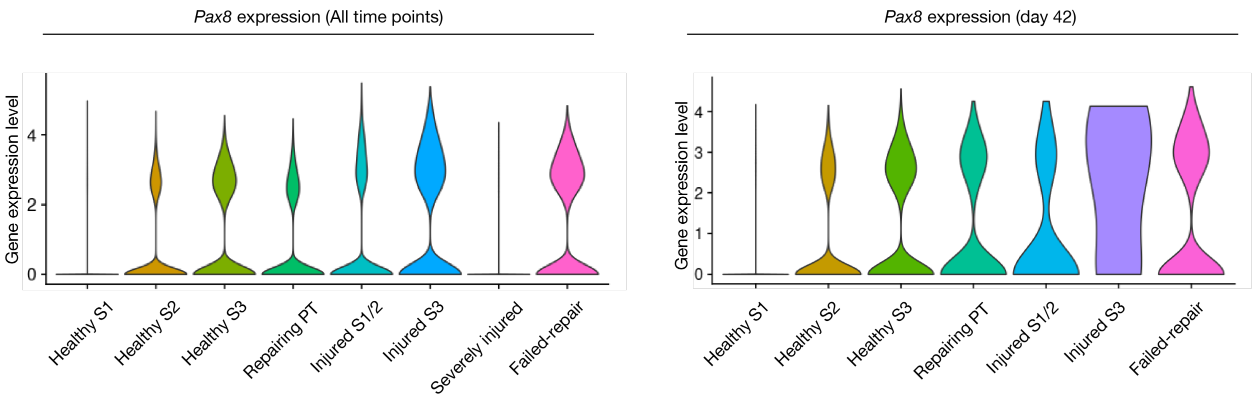


**Fig. S7. *Pax8* expression among proximal tubular cell subtypes**

Violin plot displaying *Pax8* expression levels among PT subtypes in mouse kidneys. All time points (left) or day 42 following IRI (right). *Pax8* gene is relatively highly expressed in FR-PTC at day 42.

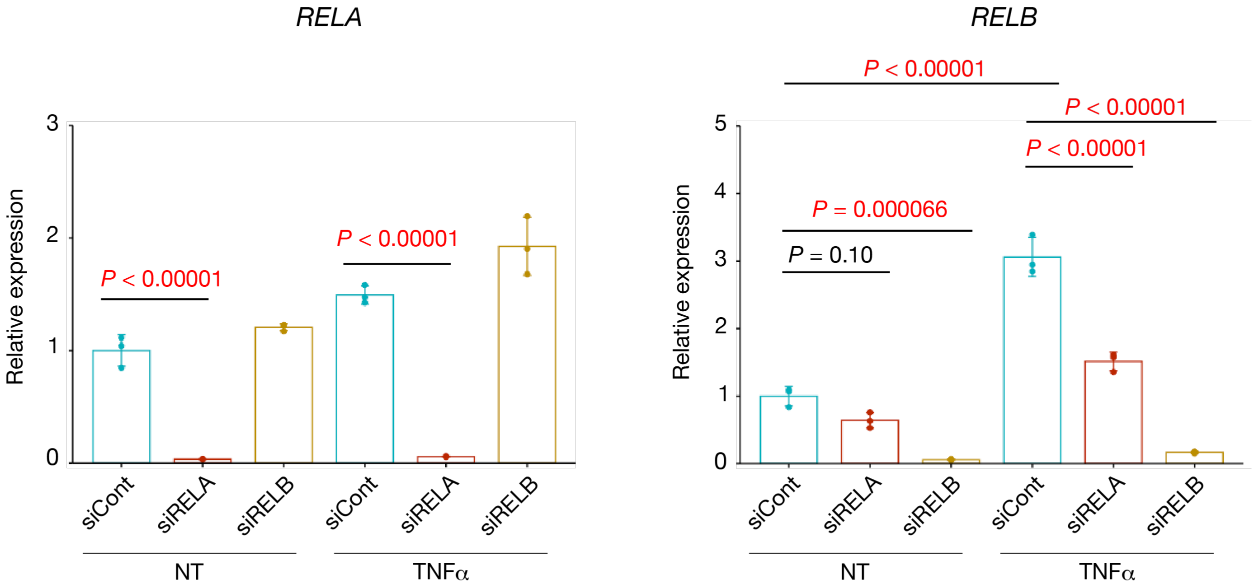


**Fig. S8. Knockdown efficiency for targeting *RELA* or *RELB* in human primary PT cells**
Quantitative PCR for *RELA* (left) and *RELB* (right) expression in primary human PTC with siRNA knockdown of *RELA* or *RELB* treated with or without TNFα (100 ng/ml). NT, no treatment. n = 3 biological replicates. Bar graphs represent the mean and error bars are the s.d. One-way ANOVA with post hoc Turkey test.

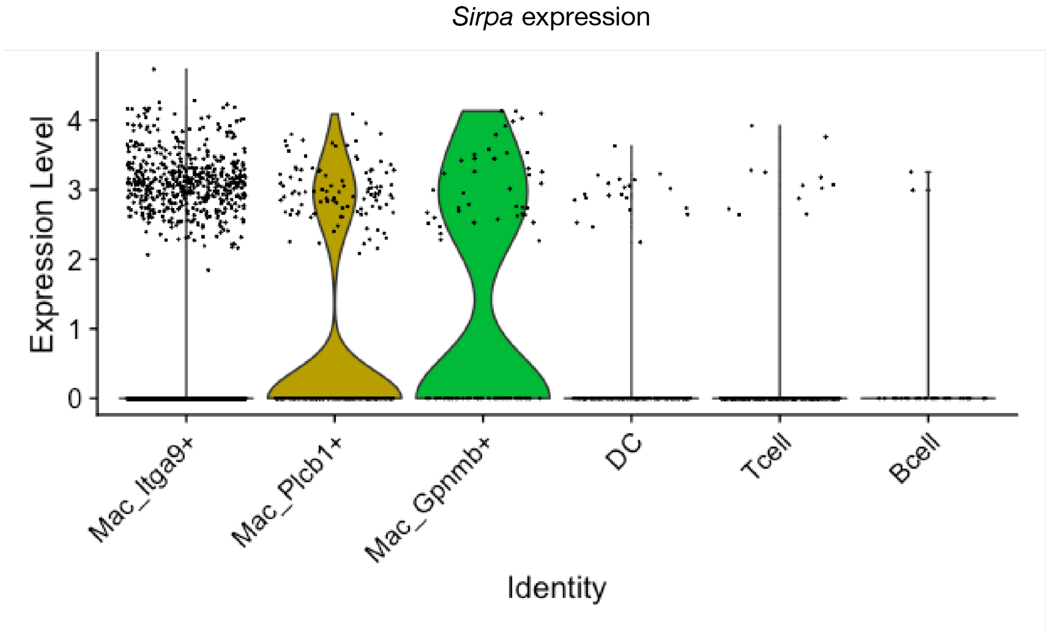


**Fig. S9. *Sirpa* expression among immune cell subtypes**
Violin plot displaying *Sirpa* expression levels among immune cell subtypes in mouse kidneys.

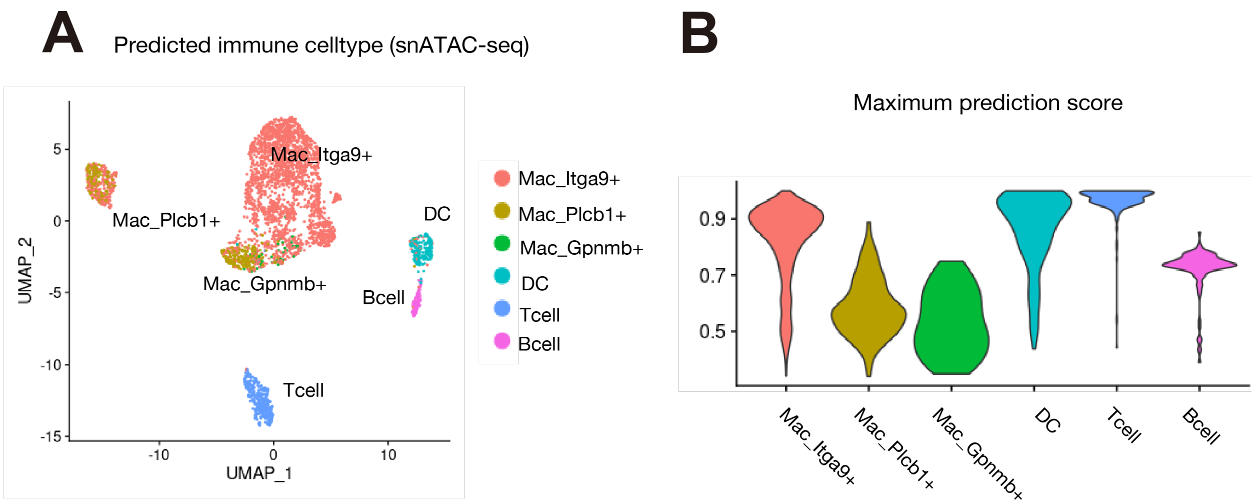


**Fig. S10. Prediction of immune cell types in snATAC-seq data with label transfer**
(**A**) UMAP plot for snATAC-seq immune cell subclustering with cell type annotation predicted with label transfer using snRNA-seq immune cell subclustering (Fig. 5A). (**B**) Violin plot showing prediction score (confidence score) for each predicted cell type.

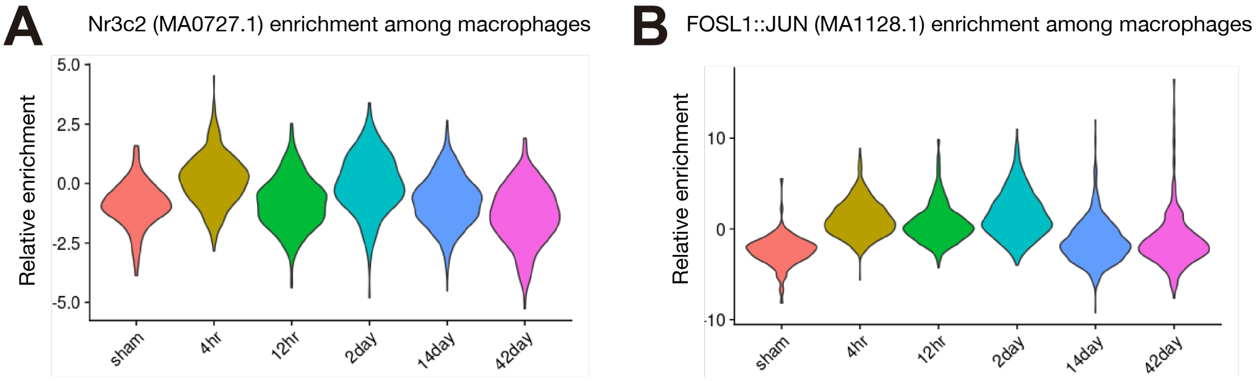


**Fig. S11. Changes in transcription factor binding motif enrichment along time course**
Violin plot displaying transcription factor motif enrichment among time points for Nr3c2 (MA0727.1) (**A**) and FOSL1::JUN (MA1128.1) (**B**) in macrophages.

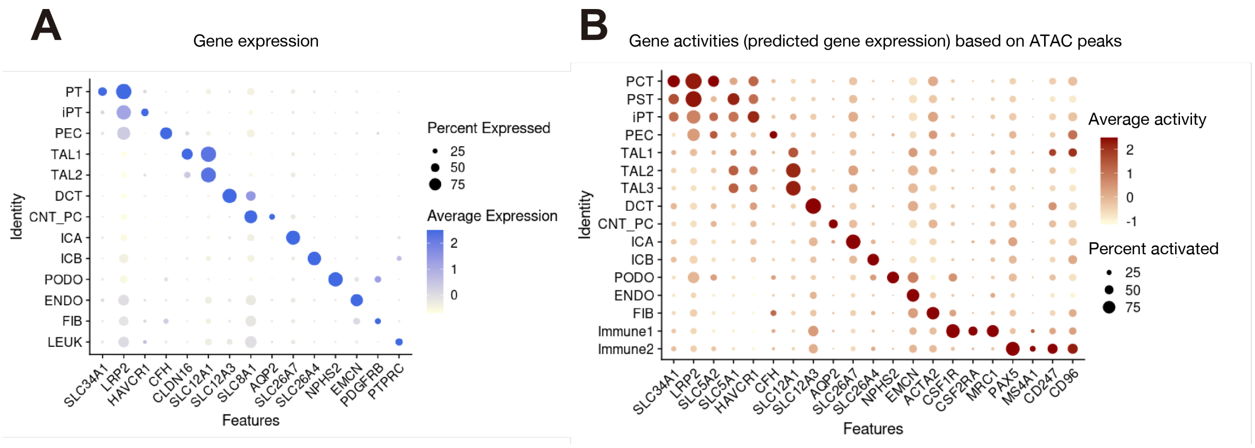


**Fig. S12. Gene expression and activity patterns for human AKI-control integrated dataset**
(A) Dot plot showing gene expression patterns of cluster-enriched markers in human kidney snRNA-seq dataset. The diameter of the dot corresponds to the proportion of cells expressing the indicated gene and the density of the dot corresponds to average expression relative to all cell types. (B) Dot plot showing gene activity patterns of cluster-enriched markers in human kidney snATAC-seq dataset

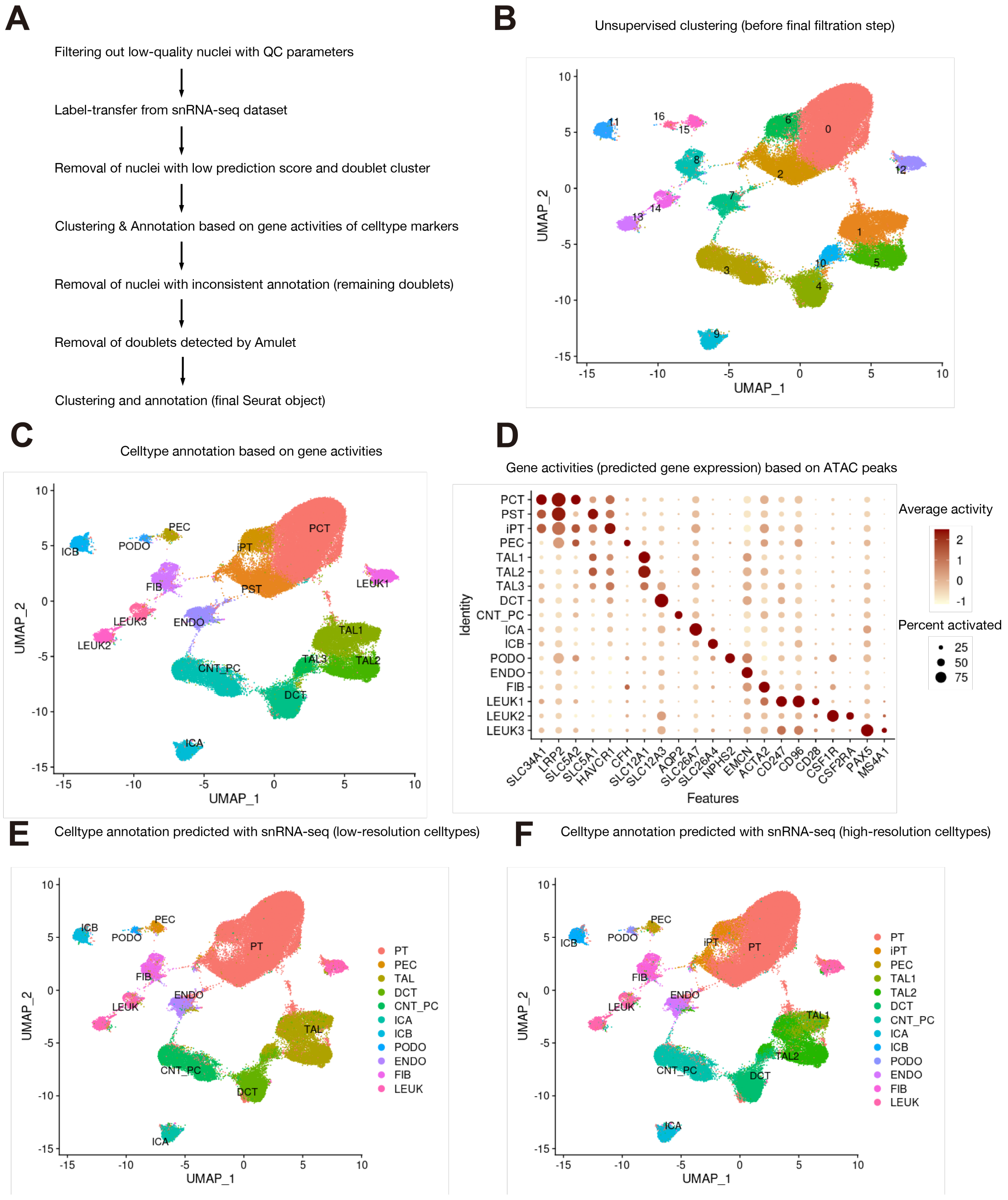


**Fig. S13. Preprocessing strategy for human AKI snATAC-seq atlas**
(**A**) Overview of preprocessing strategy. (**B**) UMAP plot of snATAC-seq dataset before the final filtering-out steps to remove the remaining doublets. (**C**) UMAP plot with cell-type annotations based on gene activities. (**D**) Dot plot showing gene activities of cell-type marker genes for (C). (**E, F**) UMAP plot with cell-type annotation prediction using human kidney snRNA-seq dataset (Fig. 6A) with low-resolution cell types (E) or high-resolution cell types (F). The nuclei with inconsistent annotations between gene activity-based (B) and low-resolution cell type prediction (E) were removed as remaining doublets, and then the doublets predicted by AMULET were further removed (A). See also Method.

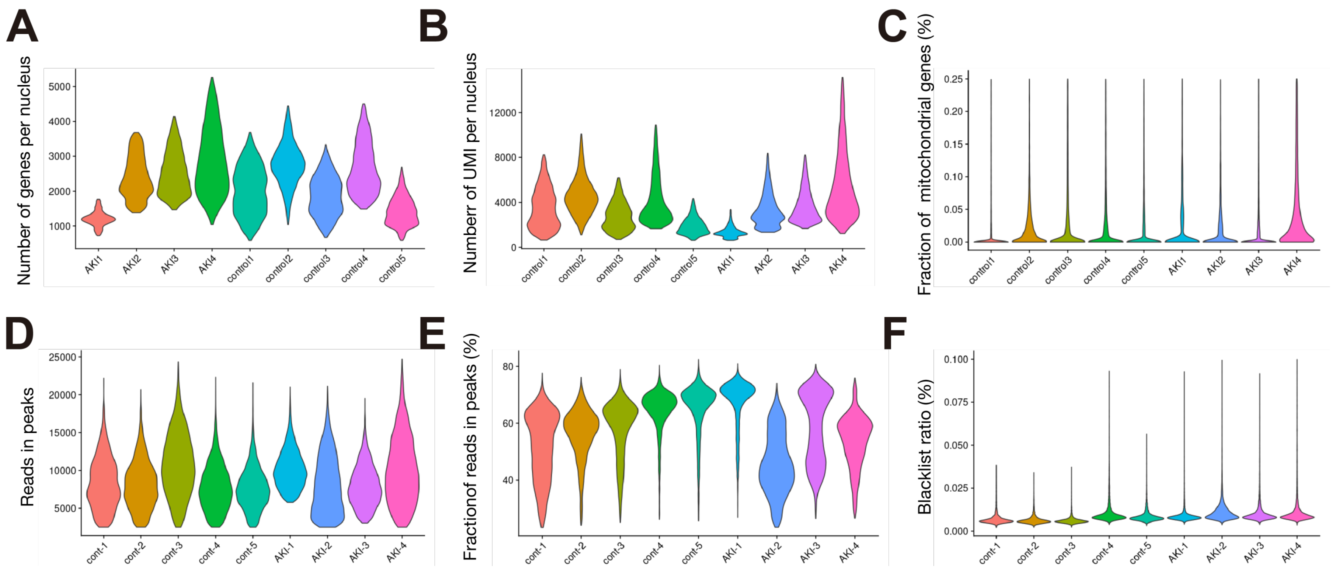


**Fig. S14. QC metrics for human kidney snRNA-seq / snATAC-seq dataset**
Violin plots showing: (**A**) number of genes per nucleus, (**B**) number of UMIs per nucleus and (**C**) fraction of mitochondrial genes per nucleus in human kidney snRNA-seq data. Violin plots showing: (**D**) number of reads in peaks per nucleus, (**E**) fraction of reads in peaks and (**F**) ratio of reads in genomic blacklist region per nucleus in human kidney snATAC-seq data.

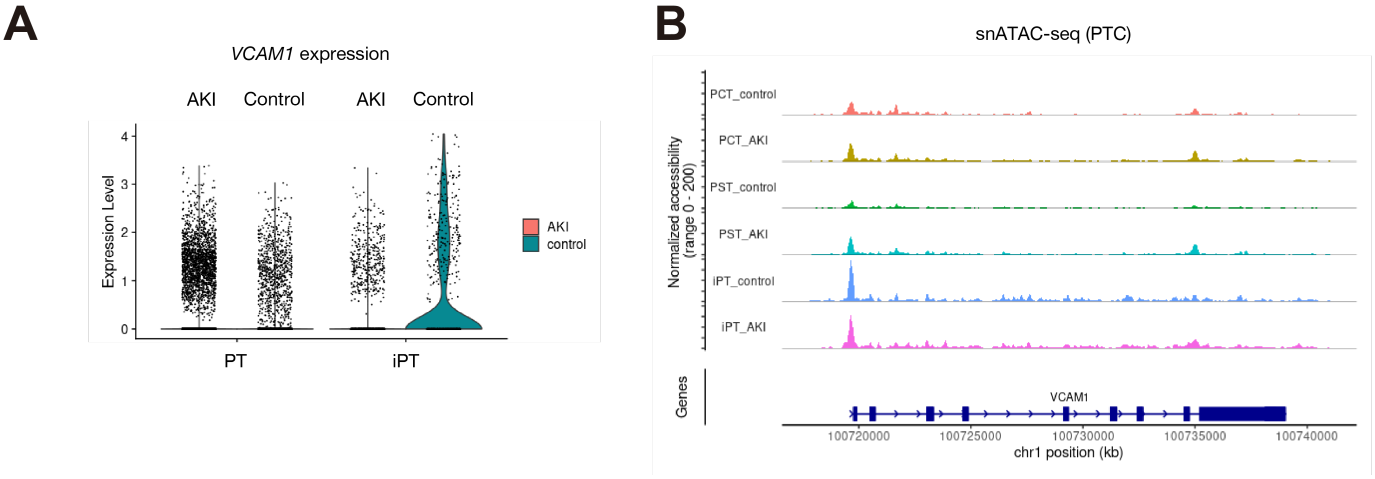


**Fig. S15. Gene expression and chromatin accessibility of *VCAM1* gene in human AKI PT**
(**A**) Violin plot displaying *VCAM1* expression levels among PT subtypes in human AKI kidneys. (**B**) Fragment coverage (frequency of Tn5 insertion) on the *VCAM1* gene in each PT subtypes of human kidneys with AKI or control.

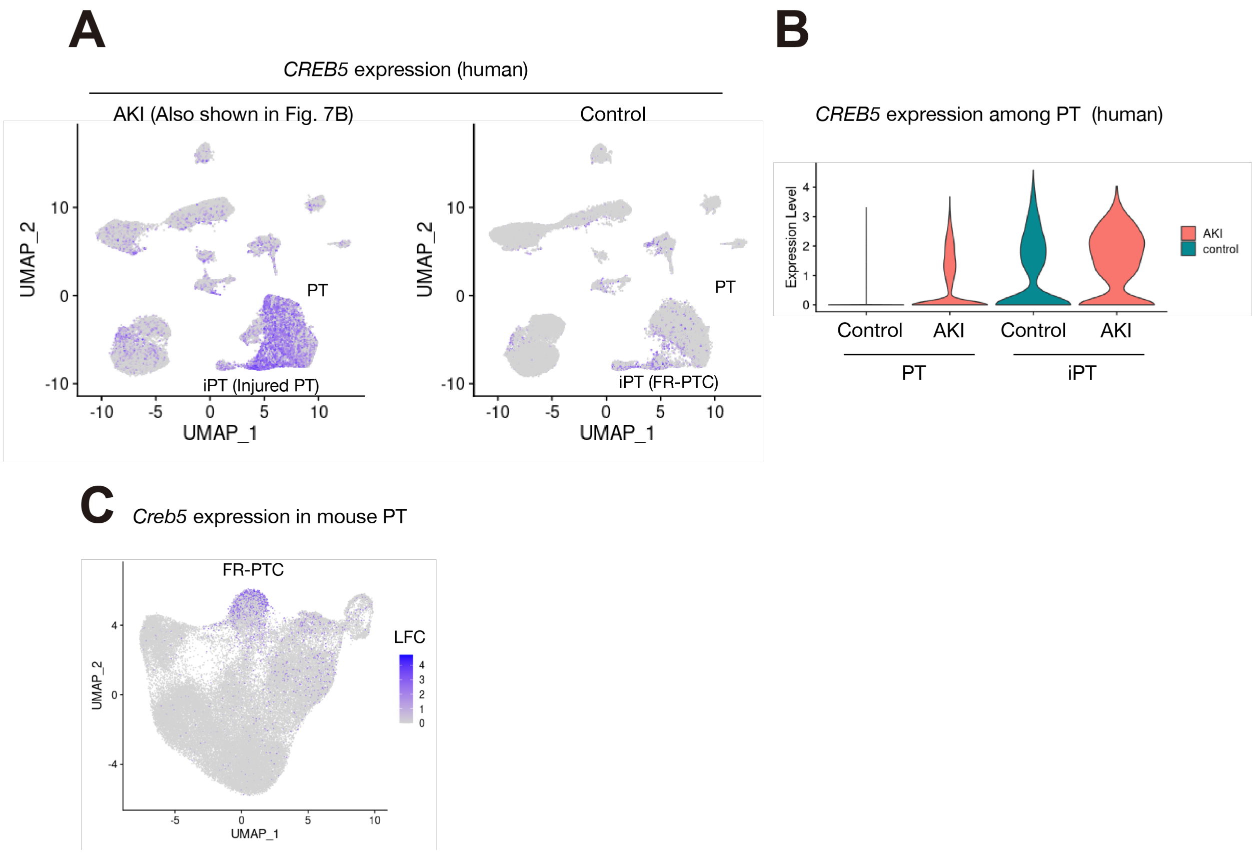


**Fig. S16. *CREB5* expression in human and mouse proximal tubular cells**
(**A**) UMAP plot showing *CREB5* gene expression in human AKI (left, also shown in Fig. 7B) or control (right) kidneys. (**B**) Violin plot displaying *CREB5* expression level among PT subtypes in human kidneys. (**C**) UMAP plot showing *Creb5* gene expression among PT subtypes of mouse kidneys with IRI. The color scale for the UMAP plot represents a normalized log-fold-change (LFC).

snRNA-seq for AKI samples (n=4)

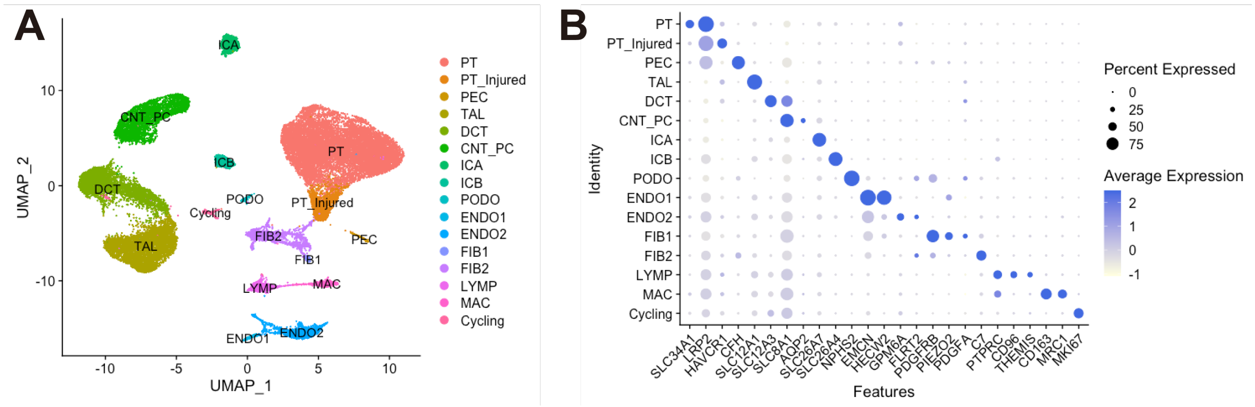


**Fig. S17. snRNA-seq aggregated dataset for only 4 AKI samples**
(**A**) UMAP plot of human AKI snRNA-seq dataset before integration with control dataset. (**B**) Dot plot showing gene expression patterns of cell type marker genes in human AKI dataset. See also Method.

| Sample ID | Gender | Age | Cr. at admission | Max Cr. | Cr. at sampling | COD |
|---|---|---|---|---|---|---|
| AKI1 | Male | 56 | 1.4 | 5.5 | 5.5 | CVD |
| AKI2 | Female | 69 | 1.2 | 5.3 | 5.3 | CVD |
| AKI3 | Female | 60 | 1.0 | 5.1 | 1.9 | CVD |
| AKI4 | Male | 55 | 1.1 | 2.4 | 1.4 | Infection |

**Table S1. AKI Patient clinical information abstracted from the medical record**
Serum creatinine value (Cr. [mg/dl]) at admission, maximum Cr. during the course (Max. Cr.), Cr. at the time of sample collection (Cr. at sampling) and cause of death (COD) were shown. CVD, Cardiovascular disease.

| **Sequences for quantitative PCR primers** | | |
|---|---|---|
| Gene | Forward primer | Reverse primer |
| *GAPDH* | GACAGTCAGCCGCATCTTCT | GCGCCCAATACGACCAAATC |
| *CCL2* | TCGCCTCCAGCATGAAAGTC | GGCATTGATTGCATCTGGC |
| *CSF1* | TGGCGAGCAGGAGTATCAC | AGGTCTCCATCTGACTGTCAAT |
| *CD47* | AGAAGGTGAAACGATCATCGAGC | CTCATCCATACCACCGGATCT |
| *RELA* | CCCACGAGCTTGTAGGAAAGG | GGATTCCCAGGTTCTGGAAAC |
| *RELB* | CAGCCTCGTGGGGAAAGAC | GCCCAGGTTGTTAAAACTGTGC |
| *CREB5* | CCCTGCCCAACCCTACAATG | GGACCTTGCATCCCCATGAT |
| *PCNA* | CAGACTATGAAATGAAGTTGATGGA | CGTGCAAATTCACCAGAAGG |
| *FOXM1* | AAAGGAGAATTGTCACCTGGAG | TGGCCATGTAAGAGTAGGGT |
| *PLK1* | CACAGTTTCGAGGTGGATGT | ATCCGGAGGTAGGTCTCTTT |

**Table S2. Sequences for quantitative PCR primers**
